# Relationship between COVID-19 infection and neurodegeneration: Computational insight into interactions between the SARS-CoV-2 spike protein and the monoamine oxidase enzymes

**DOI:** 10.1101/2021.08.30.458208

**Authors:** Lucija Hok, Hrvoje Rimac, Janez Mavri, Robert Vianello

## Abstract

Although COVID-19 has been primarily associated with pneumonia, recent data show that its causative agent, the SARS-CoV-2 coronavirus, can infect many vital organs beyond the lungs, including the heart, kidneys and the brain. The literature agrees that COVID-19 is likely to have long-term mental health effects on infected individuals, which signifies a need to understand the role of the virus in the pathophysiology of brain disorders that is currently unknown and widely debated. Our docking and molecular dynamic simulations show that the affinity of the spike protein from the wild type (WT) and the South African B.1.351 (SA) variant towards the MAO enzymes is comparable to that for its ACE2 receptor. This allows for the WT/SA…MAO complex formation, which changes MAO affinities for their neurotransmitter substrates, thus consequently impacting the rates of their metabolic conversion and misbalancing their levels. Knowing that this fine regulation is strongly linked with the etiology of various brain pathologies, these results are the first to highlight the possibility that the interference with the brain MAO catalytic activity is responsible for the increased neurodegenerative illnesses following a COVID-19 infection, thus placing a neurobiological link between these two conditions in the spotlight. Since the obtained insight suggests that a more contagious SA variant causes even larger disturbances, and with new and more problematic strains likely emerging in the near future, we firmly advise that the presented prospect of the SARS-CoV-2 induced neurological complications should not be ignored, but rather requires further clinical investigations to achieve an early diagnosis and timely therapeutic interventions.

**TABLE OF CONTENTS ENTRY:** 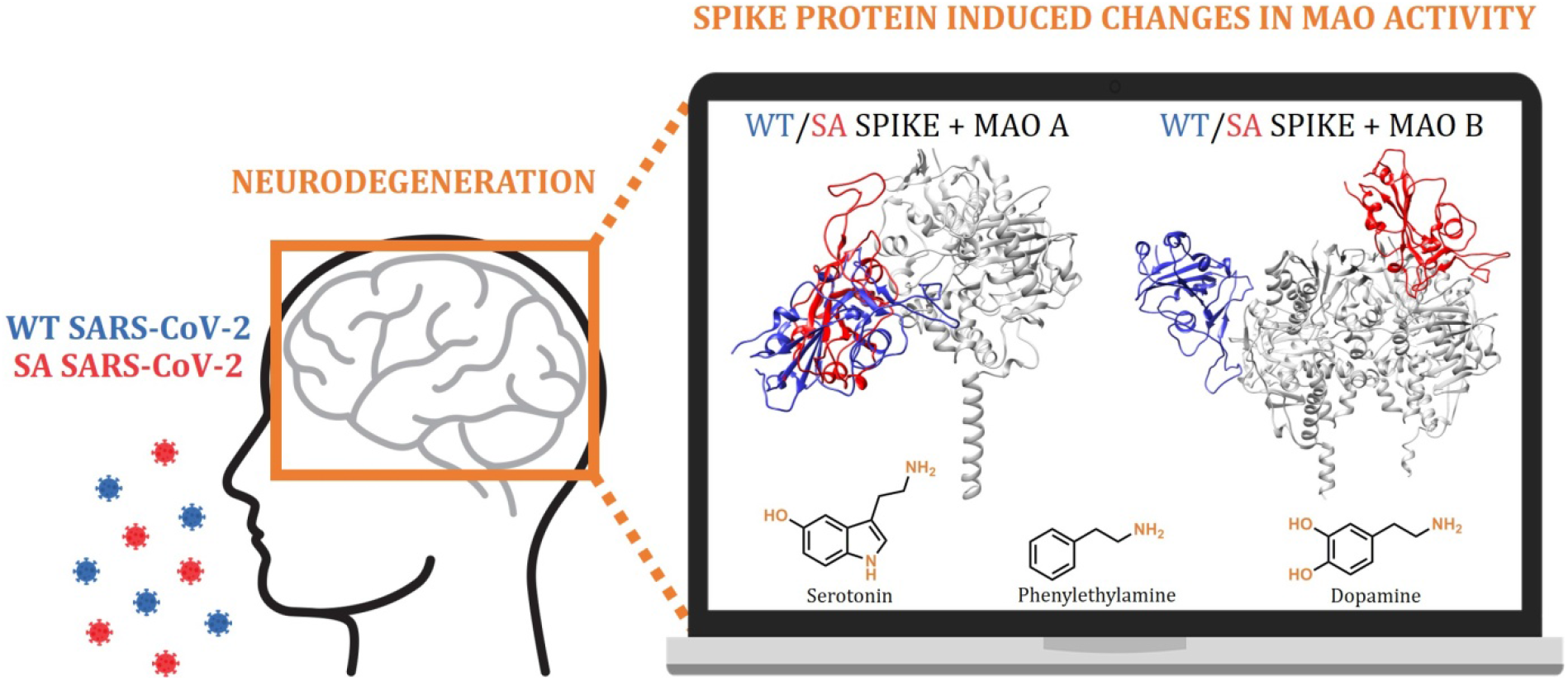

Docking and molecular dynamic simulations highlight the possibility that the interference with the brain monoamine oxidase (MAO) catalytic activity is responsible for the increased neurodegenerative illnesses following a COVID-19 infection.

## INTRODUCTION

In December 2019, a novel SARS-CoV-2 coronavirus emerged from China and spread worldwide as a pandemic, causing a public health emergency and killing over 2 million people in the first year,[1] while totalling over 4.5 million fatal outcomes by now (August 2021).[2] This infection is responsible for heterogeneous clinical disturbances, leading to severe pneumonia and the acute respiratory distress syndrome, termed COVID-19, which manifests not only as a respiratory illness but also impacts the cardiovascular, renal, and the nervous system functions.[3] Until now, this outbreak has been accompanied by a high burden on a lot of social, economic and political distress throughout the world[4] due to governmental measures of containment such as quarantine, social distancing, and lockdown. Importantly, the long-term consequences of the virus, including its effects on mental and physical health, however, might even pose a much more serious threat in the years to come.

Despite the fact that coronaviruses have not yet been linked with particular long-term neurological sequels, the occurrence of these manifestations in COVID-19 patients is becoming increasingly reported.[5– 7] Although this suggests a possibly acute or a subacute neuropathogenicity of the virus, the risk of neurological complications in patients affected by the SARS-CoV-2 is still not entirely clarified,[6,8,9] and should not be ignored.

The SARS-CoV-2 is a novel virus and its pathophysiological mechanisms in various physiological systems are yet to be fully understood. However, a lot can be learnt from the other coronavirus subtypes known to infect humans.[8] A great structural similarity between the SARS-CoV-2 and beta coronaviruses suggests a hypothesis that the SARS-CoV-2 also possesses similar neurotrophic and neuroinvasive properties. Additionally, the SARS-CoV and the SARS-CoV-2 share around 80% genome similarity[10] and use the same ACE2 host receptor to infiltrate human cells.[11] Apart from this role of the ACE2 receptor, gene expression studies have revealed that the *ACE2* gene shows the most significant co-expression and co-regulation with the aromatic *L*-amino acid decarboxylase, which is responsible for the biosynthesis of biogenic amines and the conversion of *L*-DOPA into dopamine. This indicates that ACE2 downregulation, induced by the SARS-CoV-2 infection, might be associated with concomitant alterations in the brain amine levels, which is strongly implicated in the etiology of Alzheimer and Parkinson diseases.[12] In addition, the CT/MRI scan of COVID-19 patients showed an acute necrotizing encephalopathy, a rare encephalopathy typically associated with a viral infection of the brain tissue,[13] indicating a direct CNS infection by the SARS-CoV-2. In fact, at least four known coronaviruses (HCoV-229E, HCoV-OC43, SARS-CoV, and MERS-CoV) are able to penetrate the central nervous system,[14] and the literature agrees that the CNS infection by the SARS-CoV-2 virus may promote a development of neurodegenerative disease,[15–17] especially in people already at risk.[18] Still, a significant difference between the SARS-CoV-2 and other coronaviruses is the longer length of the spike protein sequence.[19] This disparity has been suggested to confer a higher transmissibility potential to the SARS-CoV-2, making it possible for the virus to infect humans of different races and geographical origins.[19–21] It is proposed that the virus enters the CNS through different routes, including the olfactory and trigeminal nerves, the cerebrospinal fluid, the vasculature, and the lymphatic system,[22] even without an initial lung involvement. Once the virus enters the nervous system, it can bind to the highly expressed ACE2 receptor in glial cells and neurons, and from there disseminate throughout the brain.

Clinical studies show that approximately 36% of all COVID-19 patients exhibit neurological symptoms such as stroke, headache, impaired consciousness, and paresthesia.[23] COVID-19 patients can also exhibit neurobehavioral symptoms such as euphoria, anxiety, and depression, as well as cognitive dysfunction, especially in elderly patients, which are the most susceptible to the infection.[24] Accumulated evidence confirms the SARS-CoV-2 potential to invade the CNS, however, its effects at the molecular and mechanistic levels have so far only been speculations and hypotheses. Although a COVID-19 infection certainly represents a stressful event, which, on its own, may have a role in triggering neurodegeneration,[25] in this work we used a range of computational approaches to demonstrate that the SARS-CoV-2 can trigger misbalances in the monoaminergic system by binding the monoamine oxidase enzymes, MAO A and MAO B, with affinities comparable to those for its ACE2 receptor, thus causing a significant dysregulation in the way MAOs interact with their physiological substrates. Since MAO enzymes are involved in the metabolic clearance and regulation of brain amine levels,[26] including neurotransmitters dopamine and serotonin, whose disparity is strongly linked to the etiology and course of various neurological illnesses,[27,28] such a downregulation and modified MAO activity likely represent incipient stages of neurological disturbances, which are already broadly speculated in the literature.[29–32] Importantly, a potential relationship between the MAO enzymes and a SARS-CoV-2 infection has recently been proposed by Cuperlovic-Culf, Green and co-workers,[33] who used metabolomic profiling to detect a decrease in the concentration of phenylethylamine (**PEA**) metabolites within the cerebrospinal fluid and blood of COVID-19 related patients relative to healthy individuals, a trend similarly observed with more than 200 other metabolites, involving amino acids and their derivatives.[34] Knowing that MAO B preferentially degrades **PEA** in the CNS,[26] the authors ascribed this observation to a possible interference of the spike protein with the substrate entrance to the MAO B active site, thus providing a justification to our hypothesis. Additionally, it allows us to be confident that our work aids in identifying the critical role of the MAO enzymes towards an increased incidence of neurological disorders in the SARS-CoV-2 infected individuals, therefore placing a neurobiological link between these two conditions in the spotlight.

## RESULTS AND DISCUSSION

### Interactions between the ACE2 receptor and the spike protein from the WT SARS-CoV-2 and its South African variant B.1.351

The SARS-CoV-2 infiltrates human cells through an interaction between the virus S1 spike protein and the ACE2 receptor, a mechanism that has been extensively studied and characterized using various structural[35,36] and computational[37–39] techniques. Therefore, we felt it was useful to employ our computational setup to find relevant binding poses and dynamical features of the spike protein-ACE2 complexes and benchmark the obtained results with relevant literature data. By doing so, we have considered the wild-type (WT) virus and its B.1.351 South African (SA) variant, which is known to possess a higher ACE2 binding affinity,[40] an increased transmissibility and infectivity, and more severe clinical outcomes,[40,41] all of which make it a good model to discuss relative differences among strains. Therefore, after a docking analysis had suggested relevant binding poses as starting points for the molecular dynamic simulations, the latter identified a representative structure of the WT…ACE2 complex (Figure S1) that closely matches crystal structures.[35,36] Importantly, the subsequent MM-GBSA analysis predicted a binding free energy among proteins Δ*G*_BIND_ = –46.6 kcal mol^−1^, being in excellent agreement with –46.4 kcal mol^−1^ independently reported by Yarovski,[42] Murugan[43] and their co-workers, which served as a reference. Also, a decomposition of the binding affinity on a *per-residue* basis underlined crucial residues in both proteins that are contributing the most to the binding (Table S1). Interestingly, the top 15 spike protein residues are responsible for around 78% of the total binding energy, and all belong to the receptor-binding motif (RBM) of the receptor-binding domain (RBD), in line with other reports,[35,36,44] which confirms the validity of our calculations. The only exception is Lys417 with a notable contribution of –0.95 kcal mol^−1^, which forms a salt bridge with Asp30 from ACE2, as demonstrated earlier.[35,36,42] Also, within residues disfavouring the binding, one notices that the first two residues, Asp405 (0.99 kcal mol^−1^) and Glu406 (0.80 kcal mol^−1^), do not belong to the spike protein RBM area, while the third one, Glu484 (0.66 kcal mol^−1^), is one of the residues that is mutated in the SA variant to Lys484, where it exhibits a reduced unfavourable contribution by 0.13 kcal mol^−1^, from 0.66 kcal mol^−1^ in WT to 0.53 kcal mol^−1^ in SA (Table S1). Also, a highly unfavourable contribution from Ser19 in ACE2 (+2.42 kcal mol^−1^) agrees with the reported virus ability to improve binding upon changing the nearby environment.[45]

To put these values into a context, let us discuss data for a more contagious B.1.351 SA variant, first identified in South Africa in October 2020.[46] It carries the N501Y, E484K and K417N mutations in the RBD area[47] that confer an increased antibody resistance.[48] The overlap between the binding poses (Figure S2) does not reveal any significant difference in the way the SA variant approaches ACE2 relative to the WT, yet the identification of specific residues governing the interaction shows insightful aspects. Similarly to the WT, in the SA variant all of the top contributing residues belong, without exceptions, to the RBM area. Still, to our surprise, the most dominant residue is Tyr501, which is mutated from Asn501 in WT. Besides, its individual contribution of –9.11 kcal mol^−1^ surpasses all WT residues and is solely responsible for around 17% of the total binding energy. Specifically, through the N501Y mutation, the SARS-CoV-2 increases the individual contribution of this residue by as much as –6.7 kcal mol^−1^, which is both highly significant and highly disturbing, knowing that this mutation is well conserved and present in the UK and Brazilian strains as well,[47] although most likely independently evolved.[49] The reasons for the increased Tyr501 contribution are threefold; Tyr501 forms (i) O–H…O hydrogen bonds with Asp38, (ii) cation…π interactions with Lys353, and (iii) T-shaped C–H…π contacts with Tyr41, which is, amazingly, exactly the same binding environment demonstrated in the UK variant,[50] where this is the only RBM mutation. Overall, this results in a significantly higher binding affinity of the SA strain towards ACE2, Δ*G*_BIND_ = –54.8 kcal mol^−1^, and directly translates to its higher infectiveness, strongly coupled features well-demonstrated across species.[38] At this point, it is worth to stress that Δ*G*_BIND_ values obtained by this approach are somewhat overestimated in absolute terms. This is a known limitation of the employed MM-GBSA approach, as extensively discussed in a recent review by Homeyer and Gohlke,[51] which also underlined its huge potential in predicting relative binding energies in biomolecular complexes [39,42,43,51], precisely how this approach was used and discussed here. In this context, our analysis successfully reproduced the higher affinity of the SA strain, being in excellent agreement with experimental data,[40] thus further validating our computational setup. It is also interesting to observe that, despite the three mutations in the RBD area, the order of contributing residues is mostly unchanged among strains, which underlines the significance of single point mutations within this structural element and raises concerns that further mutations might likely offer even more problematic SARS-CoV-2 variants. Overall, we can summarize that, through the three mutations (N501Y, E484K, K417N), the SA variant increases its ACE2 affinity by –5.8 kcal mol^−1^, being solely responsible for almost 70% of the overall affinity gain (–8.2 kcal mol^−1^) between the SA strain and ACE2. This strongly confirms the hypothesis that positively selected virus mutations convey benefits regarding immune evasion and viral fitness, but also for the ACE2 binding, thus contributing to the evolution rate and expectedly causing higher disturbances in the infected organisms.

### Interactions between the MAO enzymes (MAO A and MAO B) and the spike protein from the WT SARS-CoV-2 virus and its South African variant B.1.351

After establishing that the WT and SA strains recognize ACE2 in almost identical ways, mainly through their RBM units, and that the SA…ACE2 complex is linked to a higher affinity, with both aspects being firmly in line with experimental data, we proceeded by applying the same protocol for the interaction among strains and the MAO isoforms. In each case, docking analysis elucidated ten most favourable binding poses (Figure S3), which were submitted to MD simulations (for details, see Computational Details), and trajectories with the highest protein-protein affinities were used for further analysis. Elucidated representative structures are shown in Figures S4 and S5, while the calculated affinities and their decomposition on a *per-residue* basis are given in Tables S2 and S3. In addition, the overlap of the resulting spike protein binding poses to each MAO isoform is depicted in Figure 1.

**Figure 1.**
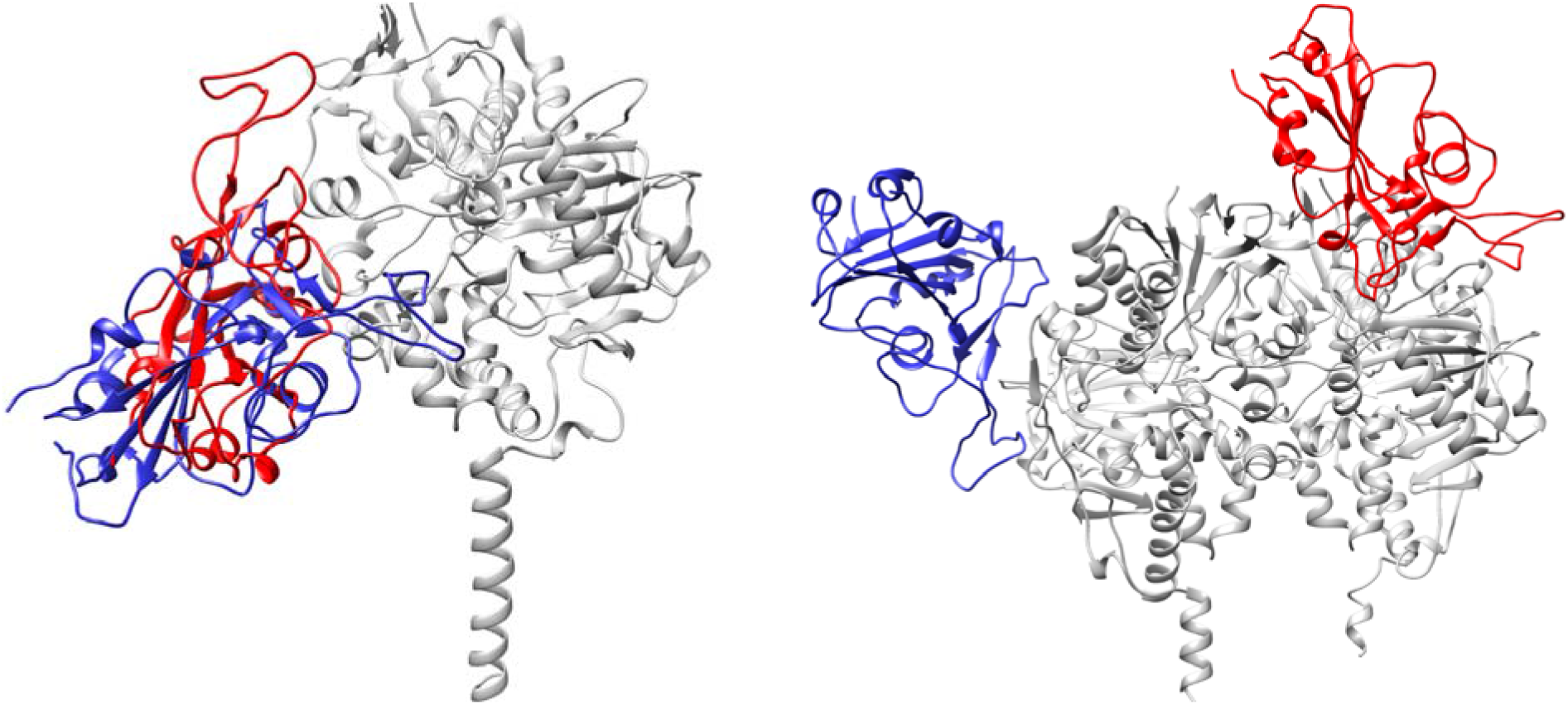
Overlap of the most favourable binding positions of the WT (in blue) and the SA (in red) spike protein in complex with the MAO enzymes (in gray), MAO A (left) and MAO B (right), as elucidated from molecular dynamic simulations.

As it was the case with ACE2, both SARS-CoV-2 variants predominantly interact with the MAO enzymes through their RBM region. This is evident from the fact that a majority of crucial interacting residues belongs to this structural element of the spike protein. This holds especially for the WT strain, while for the SA strain, it is observed that the mutated Asn417 residue becomes very significant in binding MAO A with a second largest individual contribution of –4.08 kcal mol^−1^. Interestingly, the WT binds both MAOs with an almost identical exergonicity, Δ*G*_BIND_ = –38.3 kcal mol^−1^ for MAO A and Δ*G*_BIND_ = –38.1 kcal mol^−1^ for MAO B. Given that the stability of the WT…ACE2 complex was estimated at Δ*G*_BIND_ = –46.6 kcal mol^−1^, a difference in a few kcals mol^−1^ certainly allows a formation of the matching WT…MAO complexes. In MAO A, the interaction with the WT spike protein (S) is dominated by S-Phe486, which establishes cation…π interactions with Lys357 in MAO A (Figure S4). This is followed by S-Ser477, which joins S-Thr478 in forming hydrogen bonds with Glu329, and by S-Thr500 which interacts with the side chain carbonyl group from Gln293. It is also worth mentioning that S-Lys417 forms a salt bridge with Glu159, which imitates an analogous interaction with Asp30 from the ACE2 receptor. Considering MAO B, the relative importance in spike protein residues is reversed relative to MAO A, making S-Tyr449 the most dominant residue, which is engaged in a hydrogen bonding with Gln49 and in a T-shaped C–H…π interaction with Tyr53 in MAO B (Figure S4). Interestingly, S-Glu484, which is mutated to Lys484 in the SA strain, is the first in disfavouring binding between the proteins (+1.73 kcal mol^−1^). It is placed in close vicinity of the crucial MAO B residue, Arg307, yet not interacting with it, thus its unfavourable contribution.

When a more contagious SA variant is concerned, its affinity for ACE2 is higher relative to the WT, but it is surprising that its tendency to bind both MAOs is increased as well. This is particularly interesting for MAO A, where the binding pose for the SA strain is almost identical to the one established by the WT (Figure 1), yet the affinity is increased by 10.7 kcal mol^−1^ to Δ*G*_BIND_ = –49.0 kcal mol^−1^. Recalling that its affinity for ACE2 is Δ*G*_BIND_ = –54.8 kcal mol^−1^, again only a few kcals mol^−1^ higher, opens a possibility that the matching SA…MAO A complex could be formed. There, the two crucial residues with individual contributions exceeding 4 kcal mol^−1^ are S-Leu455 and the mutated S-Asn417, which use their backbone carbonyl and side chain amide atoms, respectively, to interact with Arg297 in MAO A (Figure S5). It is worth to emphasize that all three mutations present in the SA strain are promoting MAO A binding. As mentioned, the mutated Asn417 is the second most dominant residue (–4.08 kcal mol^−1^), while its non-mutated analogue Lys417 in the WT has a significantly lower contribution (–0.39 kcal mol^−1^). Analogously, in the SA strain, Glu484 (+0.07 kcal mol^−1^) and Asn501 (+0.51 kcal mol^−1^) are replaced by Lys484 (+0.22 kcal mol^−1^) and Tyr501 (+0.07 kcal mol^−1^), respectively, from which it follows that all three mutations present in the SA variant not only promote the ACE2 binding, but also jointly promote the MAO A complex formation by –3.98 kcal mol^−1^, which is significant.

The situation with MAO B is even more interesting. The interaction energy in the SA…MAO B complex is Δ*G*_BIND_ = –62.7 kcal mol^−1^, being the highest of all, even surpassing the stability of the matching SA…ACE2 complex by –7.9 kcal mol^−1^. This suggests that the SA variant would, following the initial ACE2 binding and cell infiltration, mainly attach to MAO B, a process that is thermodynamically very favourable, and which might appear particularly troublesome for neurological conditions. This recognition is dominated by S-Arg346, which forms hydrogen bonds with the Glu232 side chain and the Ala35 backbone carbonyl, both from the subunit B of the MAO B enzyme (Figure S5). Such a positive pairing leads to a very high individual contribution from Arg346 (–6.54 kcal mol^−1^), solely contributing to around 22% of the total binding energy. One of the reasons for the high SA…MAO B binding affinity relative to the WT lies in different MAO B areas preferred by both strains (Figure 1). While the WT position is almost exclusively located on one subunit, the SA strain is most favourably located closer to the interface between the MAO B subunits, which allows both units to participate in the binding, and which might be, at least partly, responsible for the increased affinity. Although our analysis identified that a majority of crucial residues belongs to the subunit B (Table S3), the most significant residue in MAO B is Arg242 belonging to the subunit A. Its very high individual contribution (–7.35 kcal mol^−1^) comes as a result of a stable salt-bridge with S-Glu340, which was persistent during MD simulations (Figure S6). Interestingly, despite such a favourable binding to MAO B, none of the three mutated residues emerges among those dominant for an increased complex stability. Still, all three residues make notable contributions, as the introduced Asn417 (+0.06 kcal mol^−1^), Lys484 (–0.40 kcal mol^−1^) and Tyr501 (+0.08 kcal mol^−1^) surpass the initial WT residues Lys417 (+0.11 kcal mol^−1^), Glu484 (+1.73 kcal mol^−1^) and Asn501 (+0.16 kcal mol^−1^), thus increasing the binding affinity by –2.26 kcal mol^−1^.

In concluding this section, let us emphasize that the affinity of both SARS-CoV-2 variants towards the MAO isoforms is very much comparable to that for their ACE2 receptor, thus indicating a feasibility and likeliness of the WT/SA…MAO A/B complex formation. The latter is especially supported knowing that the structural comparison of the ACE2–spike protein binding region with MAO B resulted in approximately 90% structure overlap leading to a possibility of a MAO B activity change in COVID-19 patients, despite only 51% structural similarity with the overall ACE2 structure.[33] Our results demonstrate that this recognition is particularly favourable in the case of SA…MAO B, where the calculated binding energy surpasses that of SA…ACE2 by –7.9 kcal mol^−1^, thus offering an interesting insight and perspective.

### Changes in the affinity of the MAO isoforms towards physiological substrates following an interaction with the SARS-CoV-2 variants

Lastly, we evaluated how the WT/SA…MAO complexes impact MAO activity through the affinity for their brain amines. In doing so, we considered phenylethylamine (**PEA**) for both MAO isoforms, in order to place our results in the context of experimental findings by Cuperlovic-Culf, Green and co-workers,[33] and more specific amine neurotransmitters serotonin (**SER**) and dopamine (**DOP**), which are typical substrates for MAO A and MAO B, respectively. The calculated affinities are given in Table 1 and compared to relevant Michaelis-Menten constants, *K*_M_. We note in passing that in the native MAO B, both subunits revealed comparable substrate affinities without any significant preference, in line with other reports,[52] so a more exergonic binding is considered, while for the MAO B…WT/SA complexes, the results for both subunits are given, while we mostly discuss those pertaining to the subunits directly interacting with the spike protein.

**Table 1.**
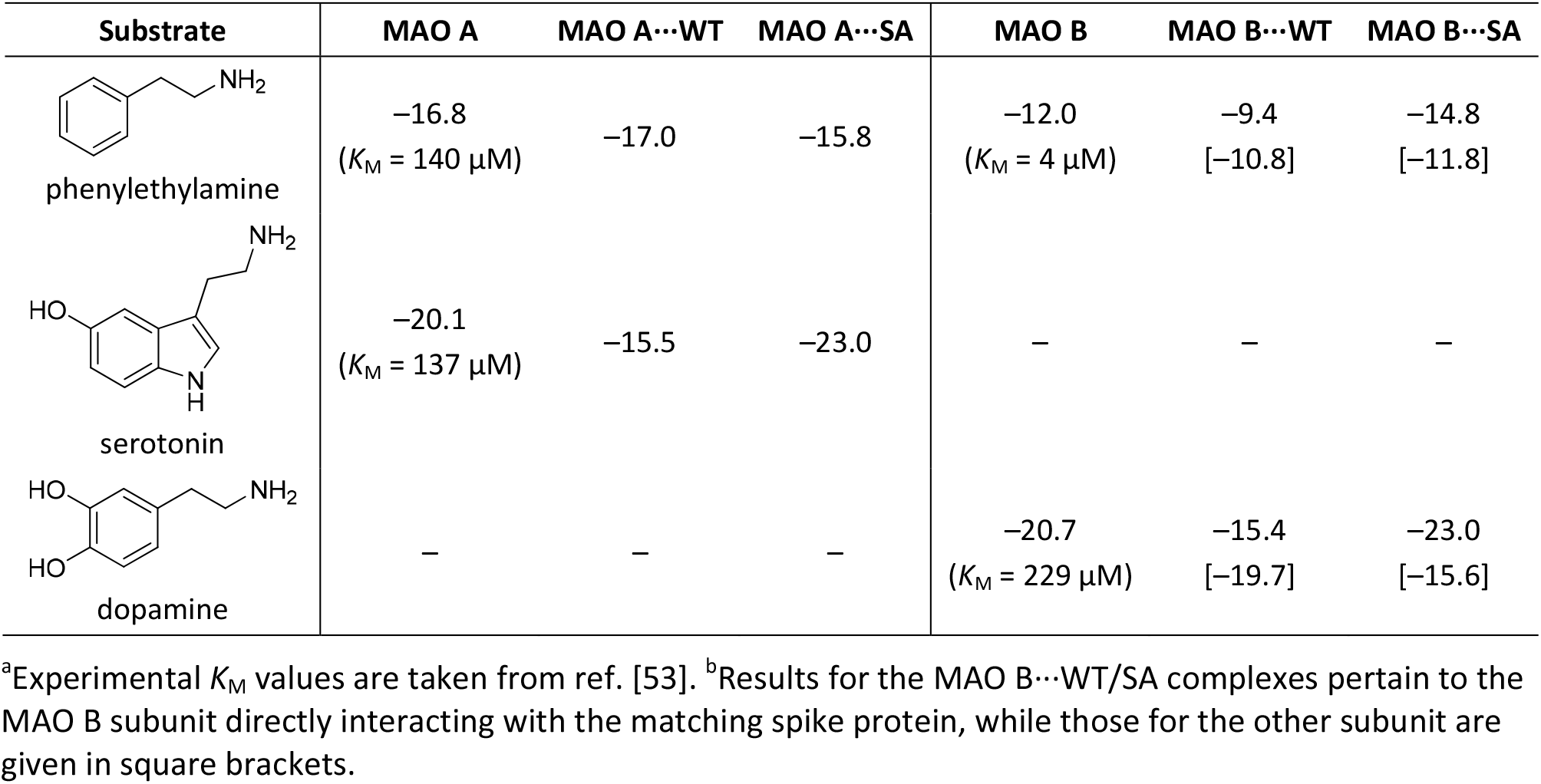
Changes in the binding affinity (Δ*G*_BIND_) between the MAO isoforms and their physiological substrates following a complex formation with the WT and SA SARS-CoV-2 variants (in kcal mol^−1^).^a,b^

| Substrate | MAO A | MAO A...WT | MAO A...SA | MAO B | MAO B...WT | MAO B...SA |
| --- | --- | --- | --- | --- | --- | --- |
| phenylethylamine | -16.8<br>($K_M = 140 \mu\text{M}$) | -17.0 | -15.8 | -12.0<br>($K_M = 4 \mu\text{M}$) | -9.4<br>[-10.8] | -14.8<br>[-11.8] |
| serotonin | -20.1<br>($K_M = 137 \mu\text{M}$) | -15.5 | -23.0 | – | – | – |
| dopamine | – | – | – | -20.7<br>($K_M = 229 \mu\text{M}$) | -15.4<br>[-19.7] | -23.0<br>[-15.6] |

<sup>a</sup>Experimental $K_M$ values are taken from ref. [53]. <sup>b</sup>Results for the MAO B...WT/SA complexes pertain to the MAO B subunit directly interacting with the matching spike protein, while those for the other subunit are given in square brackets.

The results for native MAOs show excellent agreement with the *K*_M_ data (Table 1), which lends strong credence to the employed computational setup. Specifically, **PEA** prefers binding to MAO A over MAO B, in line with the experimental affinities.[53] Additionally, the latter translate to a difference of 2.1 kcal mol^−1^, which is well-matched by our computed affinity difference of 4.8 kcal mol^−1^ in favour of MAO A. Also, **DOP** is a better substrate for MAO B than **SER** is for MAO A, again tying in with experiments. There, an even stronger agreement between sets is achieved, as the experimental affinity difference of 0.3 kcal mol^−1^ between **DOP** and **SER** is almost perfectly reproduced by the computed value of 0.6 kcal mol^−1^.

When **PEA** is concerned, the effect of the WT on its MAO A affinity is modest, being only 0.2 kcal mol^−1^ higher. Yet, for MAO B, the impact is much more pronounced, which is particularly relevant, and the observed affinity reduction following the MAO B…WT complex formation is 2.6 kcal mol^−1^. The latter indicates about two orders of magnitude lower **PEA** binding, which will inevitably lead to a lower metabolic **PEA** conversion and higher **PEA** concentrations in infected individuals. This insight strongly agrees with the mentioned experiments,[33] and helps to explain a detection of lower concentrations of **PEA** metabolites following a COVID-19 infection, thus mimicking the effects of the irreversible MAO inhibitor selegiline, whose application also increases brain **PEA** levels[54] that leads to oxidative stress.[55,56] Still, we must emphasize that our results disagree with the suggestion that the spike protein is interfering with the substrate entrance into the MAO B active site.[33] The discussed binding poses in ref. [33] were obtained through docking simulations that did not explicitly consider neither the mitochondrial membrane nor the MAO B membrane bound regions,[33] which artificially allowed the WT spike protein to reside in the area close to the membrane-mediated substrate entrances,[52] which is otherwise inaccessible and occupied by the membrane. In contrast, our simulations included entire MAO structures immersed in an explicit membrane, and after a careful inspection of the obtained binding poses in the WT…MAO A/B complexes, we found no evidence of the spike protein blocking the known substrate entrances.[52] Instead, based on our current results, we propose that the spike protein is interfering with the MAO activity by modifying the electrostatic environment in the complex, which impacts MAO affinity for their substrates.

A practically unchanged **PEA** affinity for both MAO A and the MAO A…WT complex comes as a result of an identical **PEA** binding position within MAO A in both instances (Figure2). There, its cationic amine forms a hydrogen bond with the Gln215 side chain and the carbonyl group of the FAD cofactor (Figure S7), while its ethylphenyl unit engages in a series of aromatic C–H…π and π…π stacking interactions with Tyr407, Phe352, Tyr69 and Phe208 (Table S4). Such a binding position is very much modified in MAO B, which is not surprising knowing that it is the predominant **PEA** metabolizing enzyme in the brain despite a lower affinity relative to MAO A.[53] There, **PEA** orients its aromatic unit towards FAD, and its cationic amine within three hydrogen bonds; with (i) the Tyr435 –OH group, (ii) the Gln206 side chain, and (iii) the Ile198 backbone (Figure S8). This is further modulated in the MAO B…WT complex, which strongly favours hydrophobic π…π stacking interactions with FAD and C–H…π interactions with Tyr398 and Tyr435 (Figure S9), at the expense of the hydrogen bonding contacts with –NH_3_^+^, which ultimately reduces the affinity.

**Figure 2.**
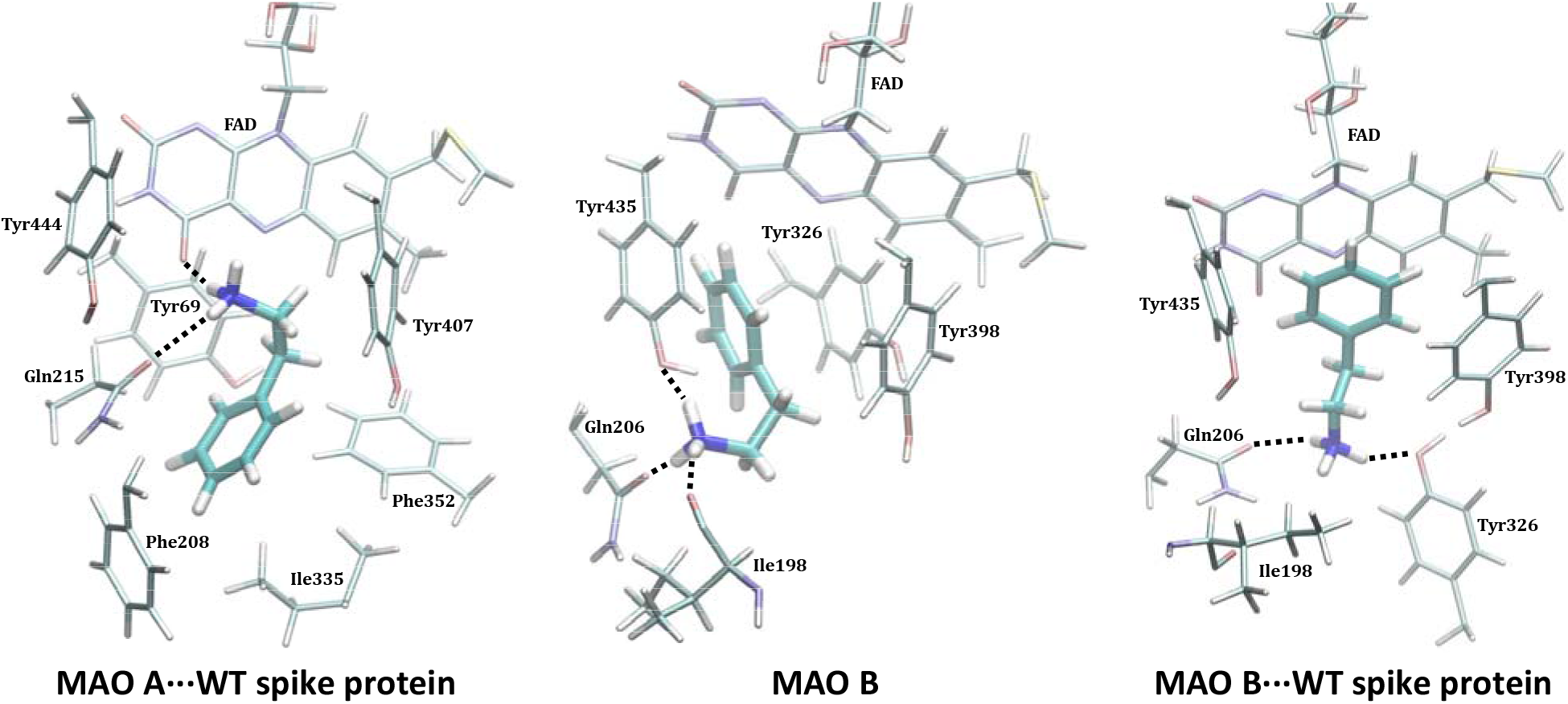
Binding position of **PEA** within the active sites of the MAO A…WT spike protein complex (left, similar in the native MAO A), native MAO B (middle) and the MAO B…WT spike protein complex (right) as obtained from MD simulations. The results for the MAO B…WT complexes pertain to the MAO B subunit directly interacting with the matching spike protein.

Encouraged by rationalizing lower **PEA** metabolite concentrations upon the WT COVID-19 infection, we continued by analyzing the more specific MAO A and MAO B substrates, **SER** and **DOP**, respectively. **SER** is a typical MAO A substrate, and its affinity, Δ*G*_BIND_ = –20.1 kcal mol^−1^, comes as a result of strong hydrogen bonding between its protonated amine and both the FAD carbonyl group and the Gln215 side chain amide (Figure S10), further supported by (i) the –OH hydrogen bonding with the Gly443 backbone amide and (ii) hydrophobic aromatic interactions with Tyr407 and Tyr444. This is significantly disrupted in MAO A…WT and results in a different binding position (Figure S11), where the mentioned three interaction motifs are replaced by the hydroxy –OH group and the protonated amine from **SER** interacting with the backbone amides of Asn181 and Ile207 (Figure S12), respectively; the former becoming the most dominant pairing in that case (Table S4). This results in a significantly lower **SER** affinity for MAO A…WT, being reduced by 4.6 kcal mol^−1^ to Δ*G*_BIND_ = –15.5 kcal mol^−1^. Such a significant impact inevitably leads to a lower **SER** metabolism upon the WT infection, which strongly corroborates experimental measurements by Shen et al.[34] On the other hand, **DOP** has the highest affinity among the studied amines, Δ*G*_BIND_ = –20.7 kcal mol^−1^, in line with its highest *K*_M_ value of 229 μM. Fascinatingly, in this case, the effect of the WT strain is also the greatest, evident in a 5.3 kcal mol^−1^ reduced affinity for MAO B. The latter is supported by a notable change in **DOP** orientation (Figure S11), during which a range of hydrogen bonding contacts in the native MAO B (Figures S13) are replaced by mostly aromatic C–H…π and π…π stacking interactions in the complex (Figure S14). Therefore, as a conclusion, somewhat higher disturbances in the dopaminergic over serotonergic pathway could be expected following the WT variant infection, which agrees with the literature.[12,30]

When a more contagious SA variant is concerned, it appears that its impact on both MAO isoforms is higher and more severe than that of the WT analogue (Figure 3, Tables S4–S5), which parallels its effect on the ACE2 receptor. Therefore, in addition to causing more disturbances in the respiratory chain, an infection with the SA strain is likely to result in more problematic outcomes for the immediate and, especially, the long-term neurological conditions. Relative to the WT, the SA strain causes the affinity of the MAO substrates to significantly increase in all cases, except for **PEA** and MAO A, where it is only slightly reduced, by 1.0 kcal mol^−1^ to –15.8 kcal mol^−1^ (Table 1). This again suggests that **PEA** and MAO A are behaving differently relative to all other instances, and that the **PEA** pathway in the affected individuals will predominantly concern the MAO B enzyme, as it was also confirmed in the WT infection.[33] There, the affinity increases by 2.8 kcal mol^−1^ to –14.8 kcal mol^−1^ (Table S5), predominantly because of favourable hydrogen bonding between its protonated amine and (i) the side chain hydroxy groups in Tyr435 and Tyr188, and (ii) the backbone amide in Cys172 (Figure S19), where the interaction with the mentioned three residues carries 56% of the total affinity. With **SER**, the effect of the SA infection is the largest and its affinity for MAO A increases by 2.9 kcal mol^−1^ to –23.0 kcal mol^−1^. The latter follows after a noteworthy change in the binding position in the SA…MAO A complex (Figure 3), which allows **SER** a range of positive and persistent hydrogen bonding contacts, including those with Gln215, Tyr444, Tyr197, Asn181 and the FAD cofactor (Figure S18), where these five MAO A residues, on their own, already contribute 17.2 kcal mol^−1^ to the binding energy, 75% in total, which is really striking. Such an affinity increase is analogously evident in **DOP**, whose affinity for MAO B becomes 2.3 kcal mol^−1^ higher and equals that for **SER** and MAO A at –23.0 kcal mol^−1^. This is again preceded by a different **DOP** binding orientation that allows it to optimize hydrogen bonding contacts with Tyr188, Ser433, Leu171 and Cys192 (Figure S20) that were all relatively insignificant for the **DOP** binding in MAO B and MAO B…WT complex, which alone are responsible for a half of the total binding energy.

**Figure 3.**
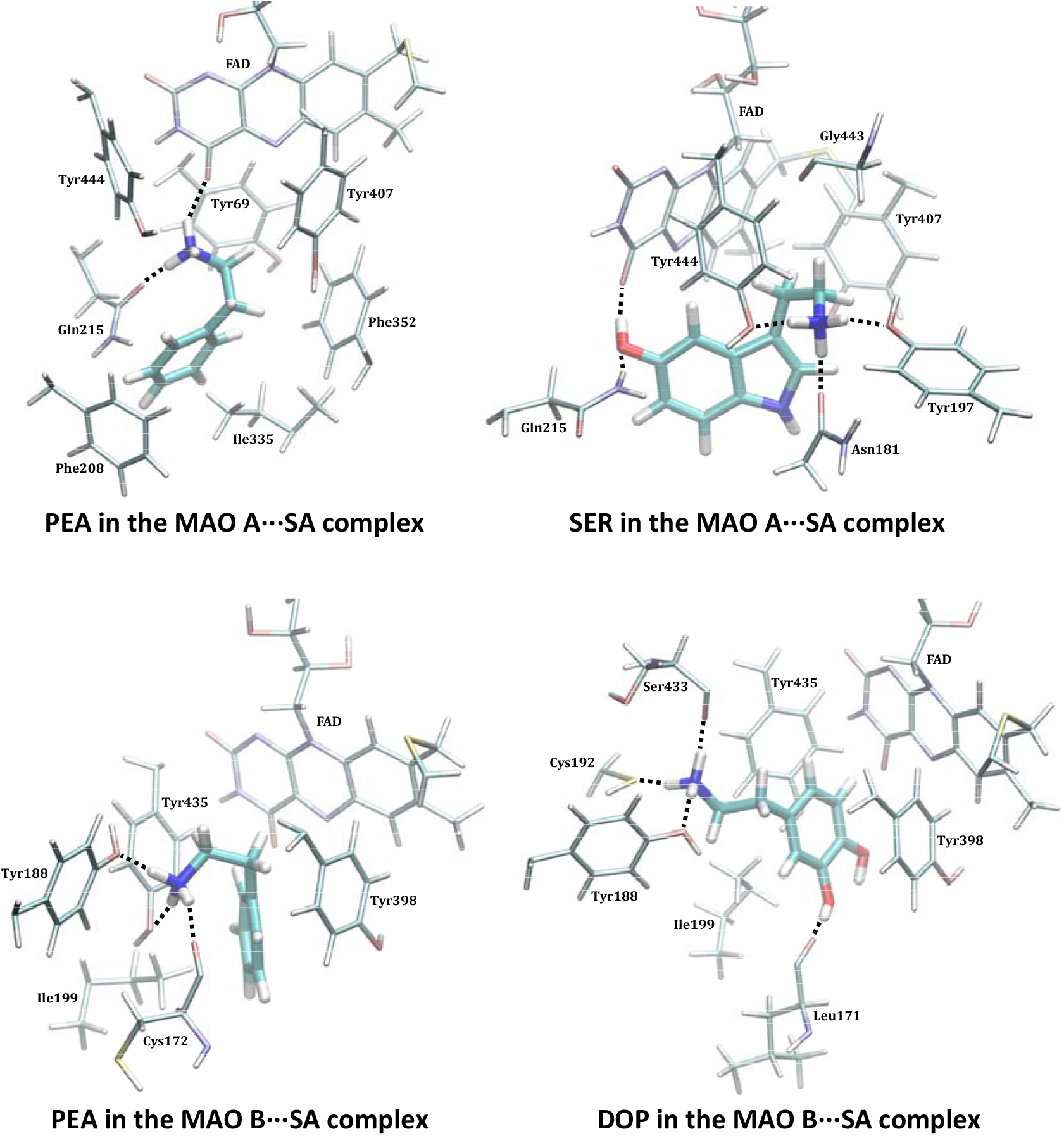
Binding positions of substrates within the active site of both MAO isoforms following the complex formation with the SARS-CoV-2 SA variant as obtained from the MD simulations. Identification of crucial interactions is presented in Figures S17–S20. The results for the MAO B…SA complexes pertain to the MAO B subunit directly interacting with the matching spike protein.

The presented results raise a serious warning that, unlike a reduction in the metabolic conversion of neurotransmitters observed in the WT, the infection with the SA mutant strain will stimulate the metabolism of the investigated brain amines, which will result in their shortage. At the same time, this will increase the production of both hydrogen peroxide and thereof derived reactive oxygen species (ROS), toxic aldehydes and ammonia, which are all by-products of the MAO-catalyzed amine degradation.[57,58] Unfortunately, all of the mentioned metabolites, along with the inflammation pathways, can induce neurodegenerative processes on their own and can further assist in their progression.

## CONCLUDING REMARKS

A combination of docking and molecular dynamic simulations reveals that the spike protein from two SARS-CoV-2 variants, namely the wild type (WT) and the mutated B.1.351 South African (SA) strain possess affinity towards the MAO enzymes that is comparable to that for its ACE2 receptor. This allows a formation of the corresponding WT/SA…MAO complexes following initial respiratory infections, with the protein-protein recognition being analogously established predominantly via residues from the receptor-binding motif in all cases. Knowing that alterations in MAO activities are a potential foundation of oxidative stress and various neuropsychiatric disorders,[27,28,57] such as Parkinson’s or Alzheimer’s disease,[59,60] the demonstrated feasibility of the WT/SA…MAO complex formation opens a possibility that the interference with the brain MAO activity is responsible for an increased development and faster progression of neurodegenerative illnesses in COVID-19 infected individuals; a disturbing medical issue that is presently widely discussed in the literature.

Our computational results show that spike protein…MAO complexes significantly lower MAO affinities towards their neurotransmitter substrates in the WT infection, thus resulting in a reduced metabolic conversion, being firmly in line with the experimentally measured trends for **PEA**[33] and a range of other metabolites in mildly affected patients.[34] However, a more severe SA variant offers even more stable complexes with both MAO isoforms, which in the case of MAO B even surpasses the stability of the matching SA…ACE2 complex. Interestingly, this leads to an increase in the MAO affinity for its substrates and, consequently, higher rates of their metabolic degradation, a trend that firmly agrees with experiments on serotonin and thereof derived conclusion that “serotonin levels would further decrease as the severity of COVID-19 increases”.[34] The latter likely promotes neurological disturbances through the immediate overproduction of hydrogen peroxide, ROS and toxic aldehydes. In this context and within the possibility for new and more contagious mutant strains likely emerging in the near future, we firmly advise that the presented prospect for the SARS-CoV-2 induced neurological complications should be carefully monitored.

It is beyond doubt that, besides changing their enzymatic function, binding of the spike protein to the MAO enzymes can additionally alter several of their roles, such as post-translational modifications or associations with protein partners.[61] This is why a possibility the SARS-CoV-2 influences MAO activity, thereby inducing neurological complications, requires further clinical investigations, which are currently scarce since most of the ongoing research focuses on drug design. Yet, our results are, to the best of our knowledge, the first in identifying a critical role of the MAO metabolic activity in this respect, therefore placing a neurobiological link between these two conditions in the spotlight and issuing a warning that it should not be ignored. In addition, we hope our work will stimulate other researchers to identify other biological systems that could be potential targets for the spike protein,[62] which could also generate various disturbances in the infected patients. Some efforts in this direction have already been made.[63]

Lastly, additional research is required to establish what effect clinically employed MAO inhibitors[64,65] might have on these pathways as, currently, there is no evidence to support either the withholding or increasing MAO inhibitors in COVID-19 treatment.

## FUNDING

This work was supported by CAT PHARMA (KK.01.1.1.04.0013), a project co-financed by the Croatian Government and the European Union through the European Regional Development Fund – the Competitiveness and Cohesion Operational Programme, and by the Slovenian Research Agency (P1-0012).

## ACKNOWLEDGMENTS

L.H. wishes to thank the Croatian Science Foundation for a doctoral stipend through the Career Development Project for Young Researchers. We thank the Zagreb University Computing Centre for computational resources on the ISABELLA cluster.

## Notes

### Competing Interest Statement

The authors have declared no competing interest.

