## Supporting Information for "Relationship between COVID-19 infection and neurodegeneration: Computational insight into interactions between the SARS-CoV-2 spike protein and the monoamine oxidase enzymes"

### TABLE OF CONTENTS

| Content | Pages |
|---|---|
| Various graphical and tabular analyses from MD simulations | S2–S26 |
| Computational details | S27–S29 |
| Equilibrated structures of MAO isoforms immersed in a bilayer DOPC membrane following MD simulations | S30 |
| References | S31–S32 |

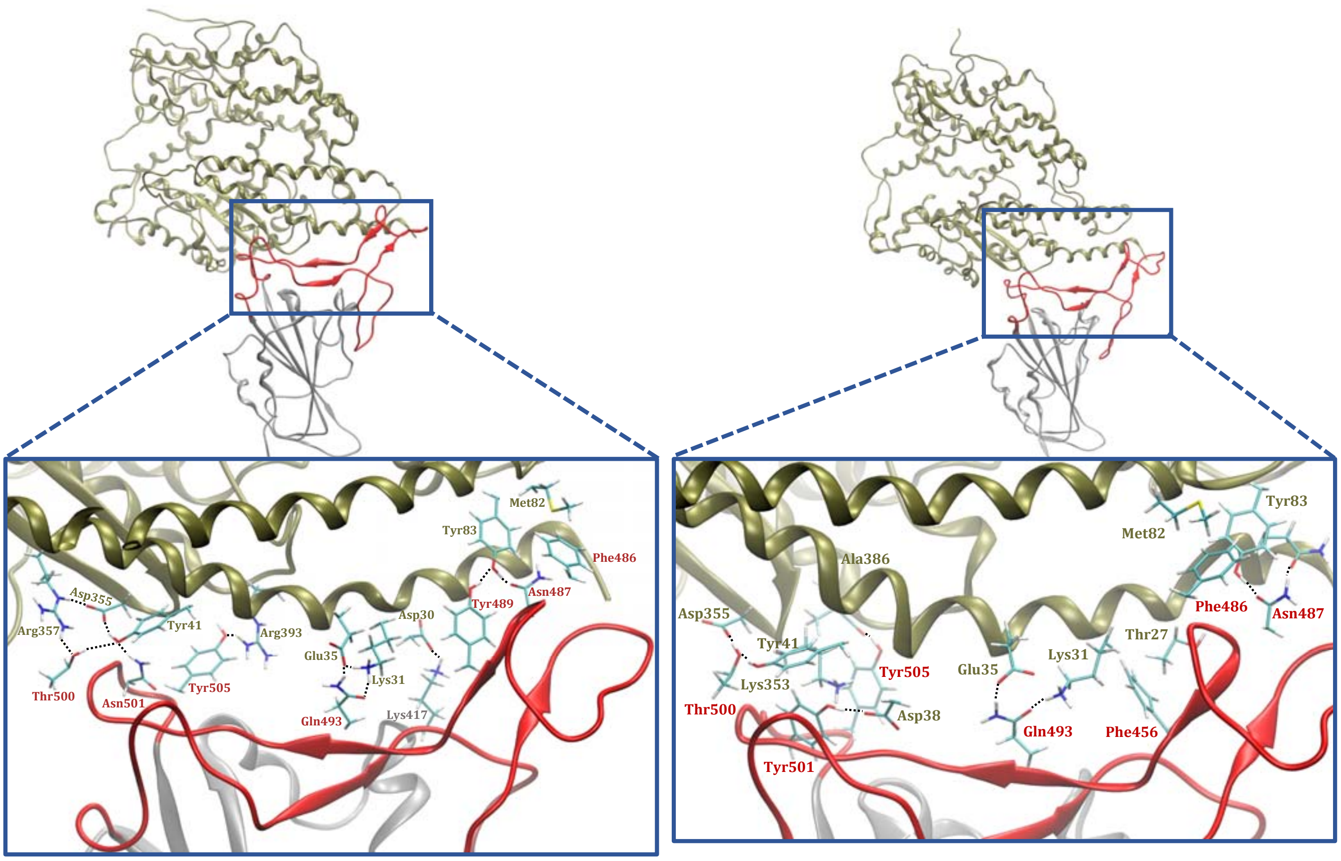


**Figure S1.** Representative structures of complexes between the **ACE2** receptor and the spike protein from different SARS-CoV-2 strains, the wild type strain (**WT**, left) and the South African strain (**SA**, right). In the overall pictures, **ACE2** is given in tan, while the spike protein is displayed in silver, with its receptor-binding motif (RBM) in the receptor-binding domain (RBD) highlighted in red. Inset figures show residues dominant for the interaction, with those belonging to **ACE2** in gold, those belonging to the spike protein RBM in red, and those not belonging to the spike protein RBM in grey.

**Table S1.** Calculated binding free energies ($\Delta G_{BIND}$) from molecular dynamic trajectories using the MM-GBSA approach, and their decomposition on a *per-residue* basis (in kcal mol$^{-1}$).[a]

| **Wild–type (WT)** $\Delta G_{BIND}$ = –46.6 kcal mol$^{-1}$ | | | | **South African (SA) strain** $\Delta G_{BIND}$ = –54.8 kcal mol$^{-1}$ | | | |
|---|---|---|---|---|---|---|---|
| **ACE2 Receptor** | | **Spike Protein** | | **ACE2 Receptor** | | **Spike Protein** | |
| Asp355 | –6.51 | Tyr505 | –6.48 | Asp355 | –6.08 | **Tyr501** | **–9.11** |
| Tyr83 | –2.43 | Gln493 | –4.20 | Tyr83 | –2.57 | Tyr505 | –6.79 |
| Tyr41 | –2.38 | Phe486 | –3.04 | Asp38 | –2.37 | Gln493 | –4.60 |
| Gln24 | –2.23 | Thr500 | –2.83 | Gln24 | –2.11 | Phe486 | –3.32 |
| Thr27 | –2.12 | **Asn501** | **–2.41** | Thr27 | –2.08 | Thr500 | –3.31 |
| Gln42 | –2.03 | Phe456 | –2.36 | Ala386 | –1.68 | Phe456 | –2.29 |
| Phe28 | –1.47 | Leu455 | –2.29 | Tyr41 | –1.65 | Leu455 | –2.22 |
| Lys31 | –1.45 | Gly496 | –2.19 | Lys353 | –1.52 | Tyr489 | –1.75 |
| Asp30 | –1.12 | Ala475 | –1.99 | Phe28 | –1.46 | Asn487 | –1.69 |
| Met82 | –1.04 | Gln498 | –1.96 | Met82 | –1.14 | Gly496 | –1.68 |
| Leu79 | –0.90 | Tyr489 | –1.75 | Leu79 | –1.02 | Ala475 | –1.43 |
| His34 | –0.88 | Asn487 | –1.63 | His34 | –1.00 | Gly502 | –1.43 |
| Leu45 | –0.67 | Gly502 | –1.44 | Lys31 | –0.93 | Gln498 | –0.83 |
| Ala386 | –0.65 | Tyr449 | –0.99 | Gly354 | –0.90 | Ser477 | –0.60 |
| Gly354 | –0.65 | **Lys417** | **–0.95** | Ala387 | –0.76 | Phe497 | –0.59 |
| Ser19 | 2.42 | Asp405 | 0.99 | Ser19 | 1.54 | Asp405 | 1.01 |
| Arg357 | 1.65 | Glu406 | 0.80 | Arg357 | 1.31 | Arg403 | 0.73 |
| Lys26 | 0.92 | **Glu484** | **0.66** | Lys26 | 0.53 | Glu406 | 0.73 |
| Asp38 | 0.52 | Gln474 | 0.64 | Gly352 | 0.45 | Gln474 | 0.55 |
| Glu37 | 0.51 | Gln506 | 0.52 | Glu37 | 0.42 | **Lys484** | **0.53** |
| Arg393 | 0.32 | Ser494 | 0.44 | Glu329 | 0.41 | Lys458 | 0.33 |
| Lys353 | 0.32 | Ser443 | 0.35 | Asn330 | 0.37 | Gln506 | 0.31 |
| Gln76 | 0.30 | Lys458 | 0.31 | Gln325 | 0.36 | Ser494 | 0.29 |
| $K_D$ = 402.5 nM[b] | | | | $K_D$ = 87.6 nM[b] | | | |

[a]Residues belonging to the receptor-binding motif (RBM) in the receptor-binding domain (RBD) of the spike protein are given in shading, while residues mutated in the **SA** strain are given in bold.
[b]Experimental dissociation constants ($K_D$) are taken from ref. [S1] and confirm a higher stability of the **SA**···**ACE2** complex over its **WT**···**ACE2** analogue.

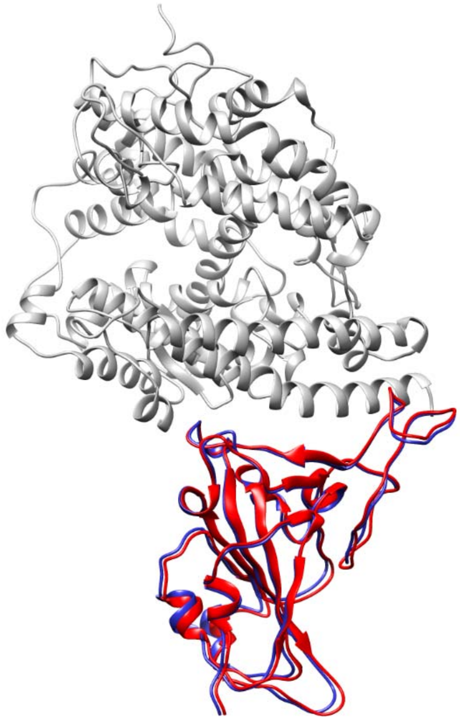

**Figure S2.** Overlap of the binding interactions between the SARS-CoV-2 **WT** (in blue) and **SA** strains (in red) with the **ACE2** receptor (in grey).

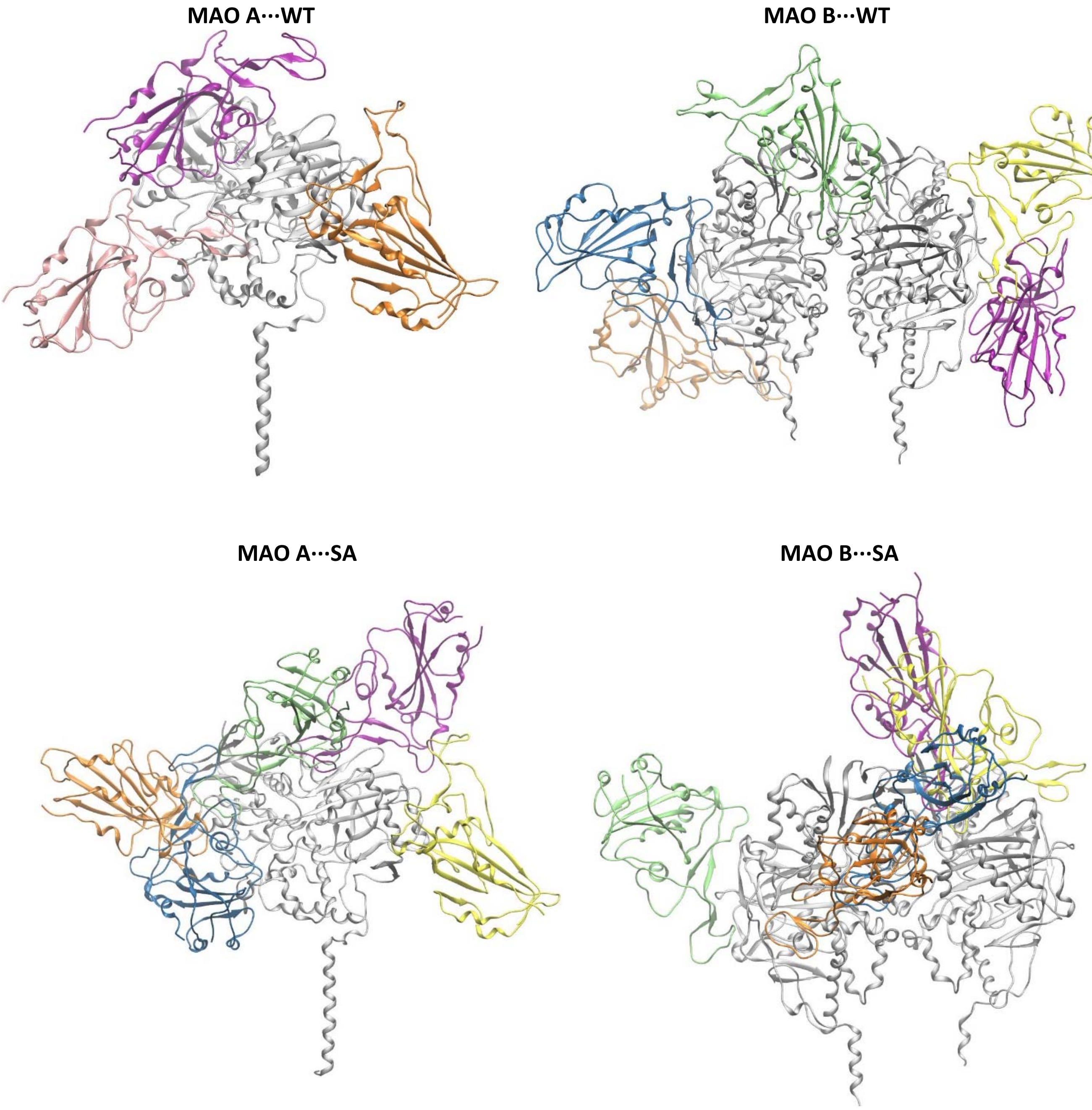


**Figure S3.** Different binding poses for the **WT** (top) and **SA** (bottom) SARS-CoV-2 spike protein in complex with the **MAO A** (left) and **MAO B** (right) enzymes obtained through docking simulations. MAO enzymes are coloured in grey, while different spike protein colours indicate different binding positions. The structure of the membrane is omitted due to clarity.

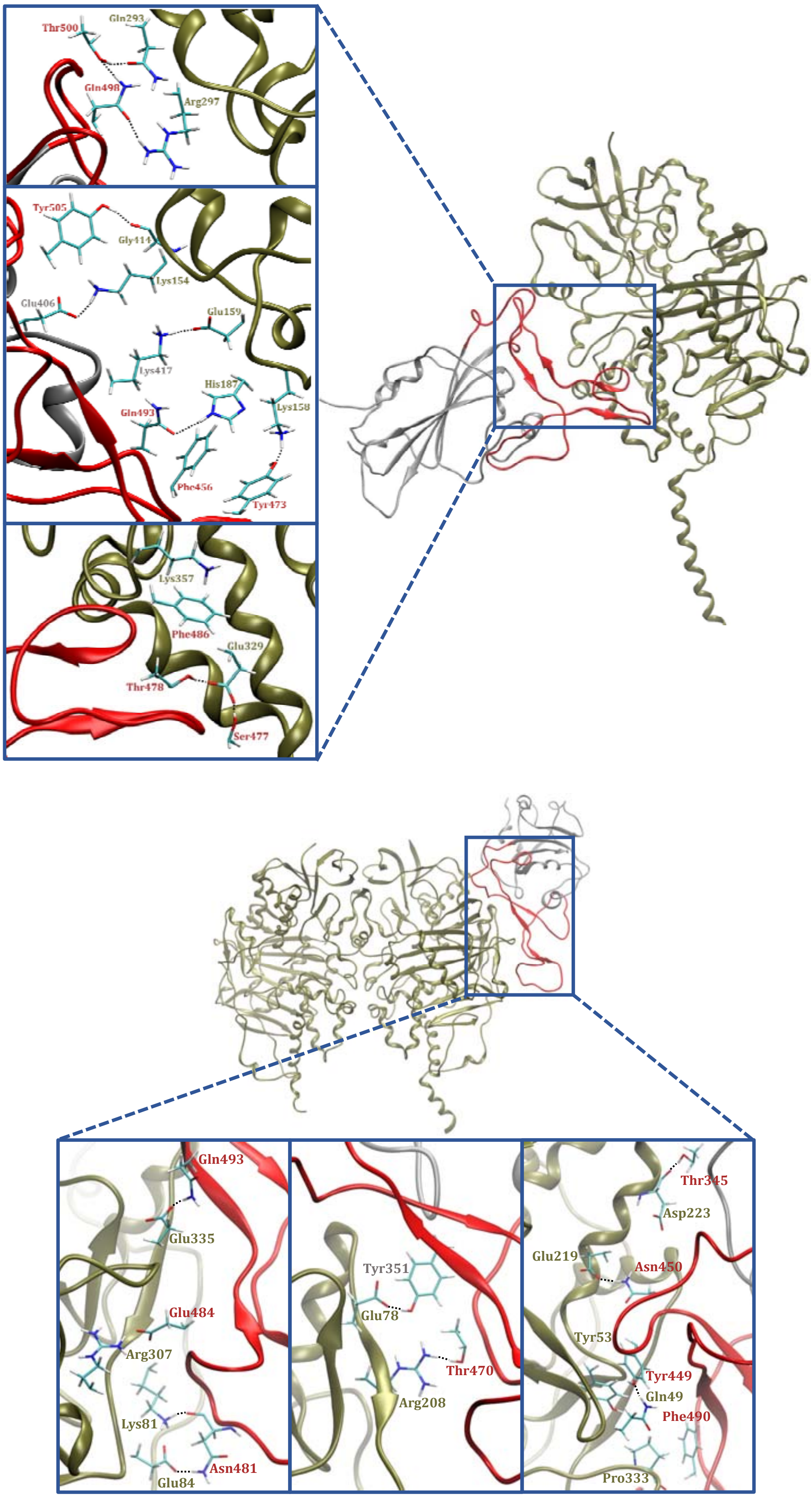


**Figure S4.** Representative structures of complexes between the SARS-CoV-2 **WT** strain and the MAO enzymes, **MAO A** (top) and **MAO B** (bottom). In the overall pictures, the MAO enzymes are given in tan, while the spike protein is displayed in silver, with its receptor-binding motif (RBM) in the receptor-binding domain (RBD) highlighted in red. Inset figures show residues dominant for the interaction, with those belonging to the MAO enzymes given in gold, those belonging to the spike protein RBM in red, and those not belonging to the spike protein RBM in grey.

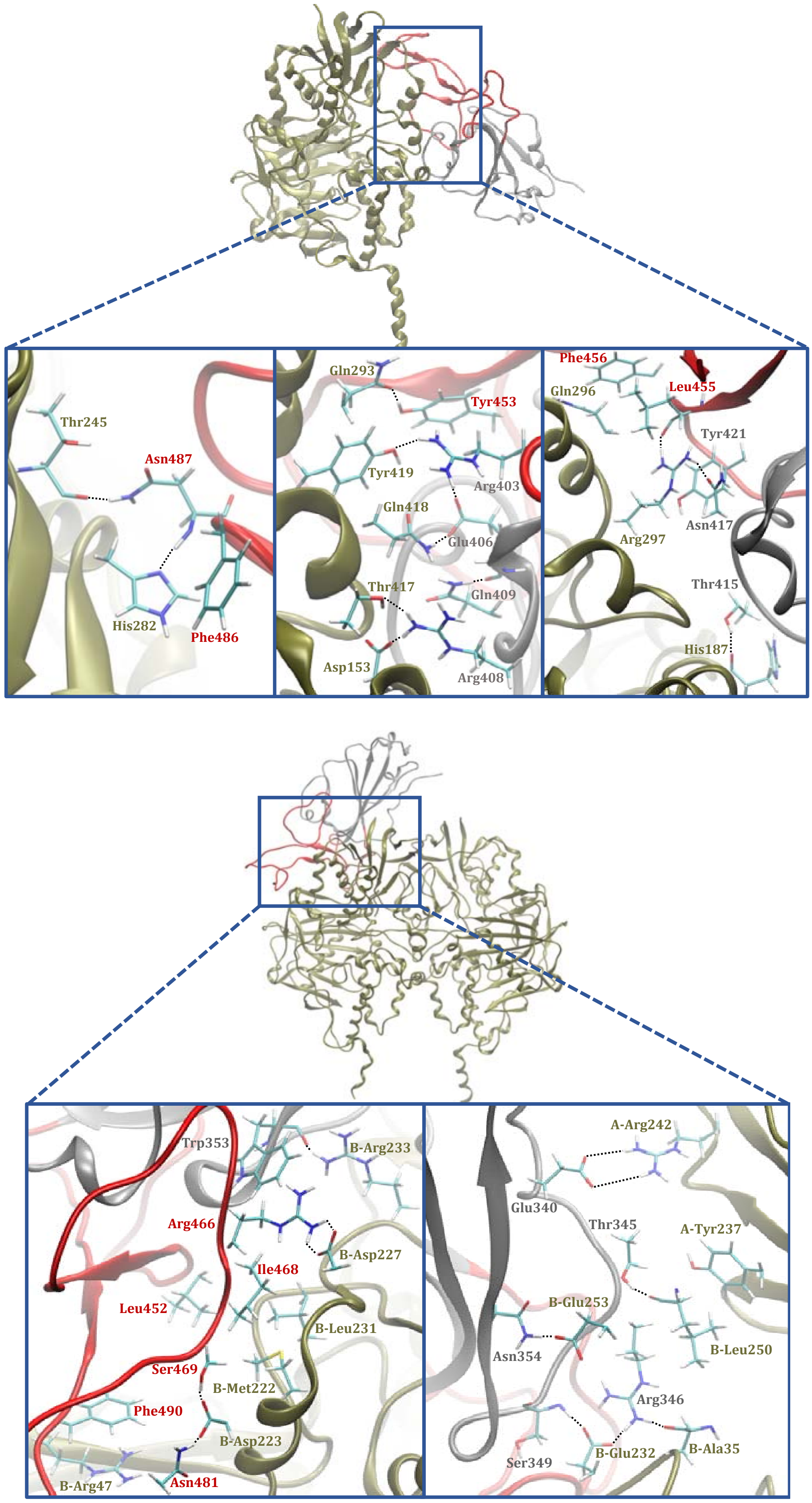


**Figure S5.** Representative structure of complexes between the SARS-CoV-2 **SA** strain and the MAO enzymes, **MAO A** (top) and **MAO B** (bottom). In the overall pictures, the MAO enzymes are given in tan, while the spike protein is displayed in silver, with its receptor-binding motif (RBM) in the receptor-binding domain (RBD) highlighted in red. Inset pictures show residues dominant for the interaction, with those belonging to the MAO enzymes given in gold, those belonging to the spike protein RBM in red, and those not belonging to the spike protein RBM in grey.

**Table S2.** Calculated binding free energies ($\Delta G_{BIND}$) from molecular dynamic trajectories using the MM-GBSA approach for the **WT** and **SA** strains with the **MAO A** enzyme, and their decomposition on a *per-residue* basis (in kcal mol$^{-1}$).[a]

| MAO A Enzyme | | | | | | | |
|---|---|---|---|---|---|---|---|
| **Wild–type (WT)** $\Delta G_{BIND}$ = –38.3 kcal mol$^{-1}$ | | | | **South African (SA) strain** $\Delta G_{BIND}$ = –49.0 kcal mol$^{-1}$ | | | |
| **MAO A** | | **Spike Protein** | | **MAO A** | | **Spike Protein** | |
| Gln293 | –3.97 | Phe486 | –5.39 | His282 | –4.12 | Leu455 | –4.76 |
| Glu329 | –3.71 | Ser477 | –5.36 | Asp153 | –3.25 | **Asn417** | **–4.08** |
| Ile415 | –1.74 | Thr478 | –4.14 | Gln418 | –3.12 | Phe486 | –3.41 |
| Glu159 | –1.69 | Thr500 | –3.31 | His187 | –2.80 | Gln409 | –2.79 |
| Arg356 | –1.58 | Tyr489 | –3.23 | Arg297 | –2.41 | Thr415 | –2.61 |
| Tyr175 | –1.47 | Tyr505 | –3.06 | Thr156 | –2.14 | Tyr421 | –2.52 |
| His187 | –1.19 | Phe456 | –2.38 | Thr417 | –2.07 | Asn487 | –2.47 |
| Gly414 | –1.12 | Gln498 | –2.14 | Thr245 | –2.05 | Phe456 | –2.10 |
| Ala289 | –0.87 | Gly476 | –1.46 | Pro413 | –1.85 | Gln414 | –2.00 |
| Lys357 | –0.84 | Tyr473 | –1.19 | Gln296 | –1.61 | Arg408 | –1.61 |
| Arg297 | –0.76 | Leu455 | –0.89 | Gly414 | –1.46 | Tyr453 | –1.54 |
| Arg172 | –0.76 | Glu406 | –0.56 | Gln293 | –1.09 | Asp405 | –1.26 |
| Pro186 | –0.68 | Phe490 | –0.54 | Tyr419 | –1.05 | Tyr489 | –1.21 |
| Leu354 | –0.54 | **Lys417** | **–0.39** | Ile415 | –1.03 | Arg454 | –1.06 |
| Lys154 | –0.47 | Gly502 | –0.35 | Pro412 | –0.91 | Gly416 | –0.55 |
| Glu188 | 1.77 | Arg403 | 1.06 | Glu159 | 0.82 | Lys424 | 1.26 |
| Lys168 | 0.62 | Gln474 | 0.63 | Glu286 | 0.51 | Asp420 | 1.21 |
| Asp330 | 0.59 | Arg408 | 0.62 | Asn292 | 0.43 | Lys378 | 0.98 |
| Arg360 | 0.45 | **Asn501** | **0.51** | Glu399 | 0.39 | Arg403 | 0.91 |
| Lys102 | 0.44 | Lys458 | 0.50 | Lys163 | 0.39 | Asn422 | 0.76 |
| Arg171 | 0.42 | Arg454 | 0.36 | Arg421 | 0.38 | Asp427 | 0.62 |
| Lys151 | 0.42 | Arg457 | 0.30 | Glu257 | 0.38 | Lys458 | 0.56 |
| Lys163 | 0.40 | Ser494 | 0.26 | Lys151 | 0.38 | Gln474 | 0.54 |

[a]Residues belonging to the receptor-binding motif (RBM) in the receptor-binding domain (RBD) of the spike protein are given in shading, while residues mutated in the **SA** strain are given in bold.

**Table S3.** Calculated binding free energies ($\Delta G_{BIND}$) from molecular dynamic trajectories using the MM-GBSA approach for the **WT** and **SA** strains with the **MAO B** enzyme, and their decomposition on a *per-residue* basis (in kcal mol$^{-1}$).[a]

| **MAO B Enzyme** | | | | | | | |
|---|---|---|---|---|---|---|---|
| **Wild–type (WT)** $\Delta G_{BIND}$ = –38.1 kcal mol$^{-1}$ | | | | **South African (SA) strain** $\Delta G_{BIND}$ = –62.7 kcal mol$^{-1}$ | | | |
| **MAO B** | | **Spike Protein** | | **MAO B** | | **Spike Protein** | |
| Arg307 | –4.04 | Tyr449 | –5.42 | Arg242 (A) | –7.35 | Arg346 | –6.54 |
| Glu78 | –2.08 | Val483 | –4.02 | Glu232 (B) | –5.27 | Ser469 | –4.12 |
| Tyr53 | –2.00 | Tyr351 | –3.84 | Asp227 (B) | –3.31 | Phe490 | –3.74 |
| Tyr80 | –1.97 | Phe490 | –3.61 | Glu253 (B) | –2.81 | Asn481 | –3.62 |
| Val211 | –1.91 | Ile468 | –2.93 | Arg220 (B) | –2.51 | Ile468 | –3.60 |
| Glu219 | –1.90 | Leu452 | –2.82 | Leu250 (B) | –2.42 | Thr345 | –3.45 |
| Leu224 | –1.73 | Thr345 | –2.61 | Leu75 (B) | –2.33 | Arg466 | –3.32 |
| Arg47 | –1.36 | Asn481 | –2.10 | Arg47 (B) | –1.91 | Ala352 | –3.32 |
| Pro333 | –1.35 | Asn450 | –2.02 | Ala35 (B) | –1.76 | Leu452 | –2.89 |
| Asp223 | –1.06 | Phe347 | –1.61 | Asn251 (B) | –1.69 | Asn354 | –2.69 |
| Leu77 | –0.80 | Gln493 | –1.47 | Leu231 (B) | –1.63 | Ser349 | –2.66 |
| Val85 | –0.79 | Tyr451 | –0.86 | Met222 (B) | –1.54 | Ala348 | –2.42 |
| Gln49 | –0.73 | Thr470 | –0.81 | Tyr237 (A) | –1.45 | Trp353 | –2.38 |
| Gly212 | –0.68 | Gly446 | –0.77 | Pro234 (B) | –1.43 | Glu340 | –2.13 |
| Lys81 | –0.51 | Arg346 | –0.72 | Arg38 (B) | –1.13 | Gly482 | –1.92 |
| Glu334 | 1.47 | **Glu484** | **1.73** | Lys230 (B) | 3.55 | Lys356 | 1.76 |
| Arg208 | 1.26 | Lys444 | 1.03 | Asn3 (B) | 1.69 | Lys444 | 1.38 |
| Lys73 | 0.80 | Glu471 | 0.65 | Arg36 (B) | 1.68 | Thr333 | 0.92 |
| Lys21 | 0.59 | Arg454 | 0.60 | Lys386 (B) | 0.98 | Glu465 | 0.74 |
| Glu74 | 0.55 | Asn354 | 0.50 | Glu219 (B) | 0.77 | Arg509 | 0.73 |
| Glu86 | 0.53 | Arg466 | 0.48 | Lys52 (B) | 0.68 | Arg454 | 0.63 |
| Lys332 | 0.48 | Val445 | 0.43 | Lys357 (B) | 0.62 | Asn448 | 0.60 |
| Arg228 | 0.41 | Ser443 | 0.39 | Glu34 (B) | 0.61 | Glu471 | 0.49 |

[a]Residues belonging to the receptor-binding motif (RBM) in the receptor-binding domain (RBD) of the spike protein are given in shading, while residues mutated in the **SA** strain are given in bold. Markings (A) and (B) denote a particular subunit of the dimeric MAO B.

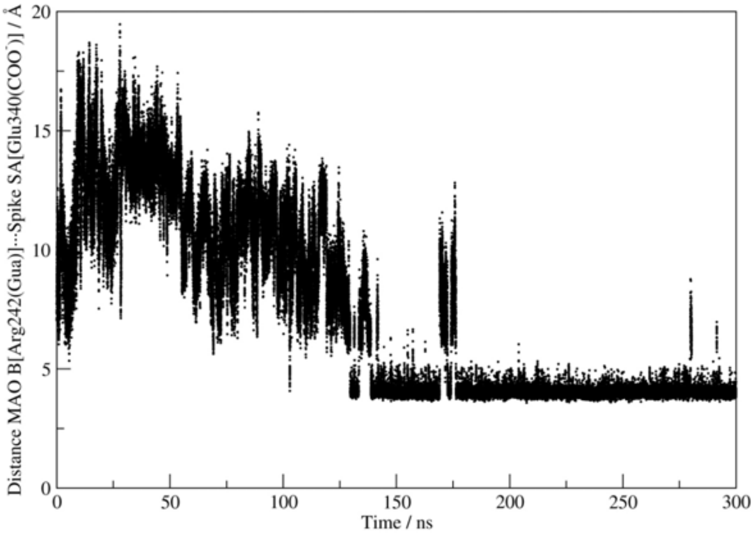


**Figure S6.** Evolution of distances between the side chain carboxylic C-atom of Glu340 in the **SA** strain and the side chain guanidinium C-atom of Arg242 from the subunit A of the dimeric **MAO B** enzyme in the **MAO B···SA complex**, indicating a stable and persistent interaction in the last 180 ns of MD simulations following the initial equilibration.

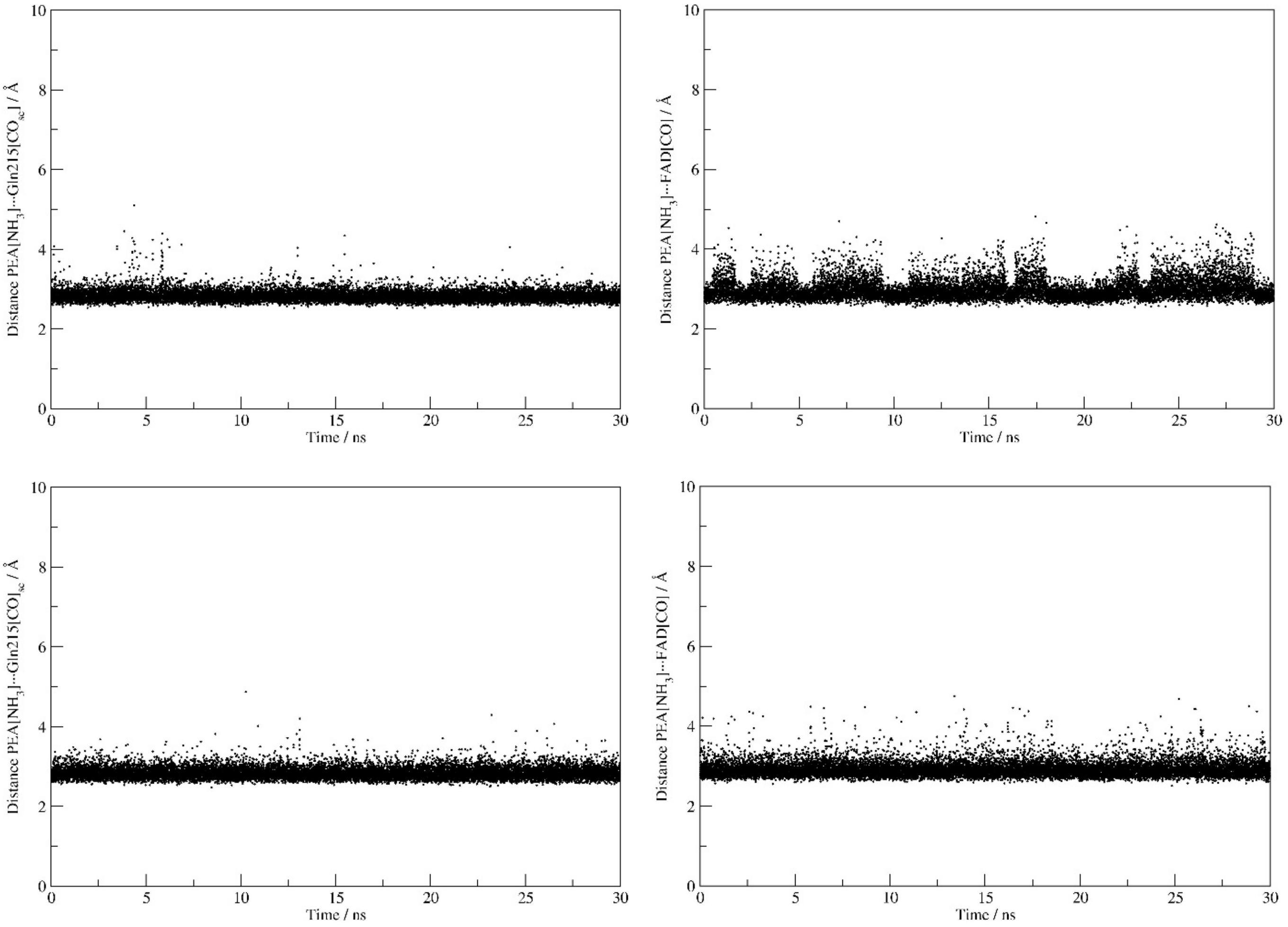


**Figure S7.** Evolution of distances between the protonated amine N-atom in **PEA** and (i) the side chain amide O-atom of Gln215 in **MAO A** (left) and (ii) the carbonyl O-atom of the FAD cofactor in **MAO A** (right). Top displays correspond to the **native MAO A**, while bottom displays pertain to the **MAO A···WT complex**, with both illustrating very similar **PEA** binding poses.

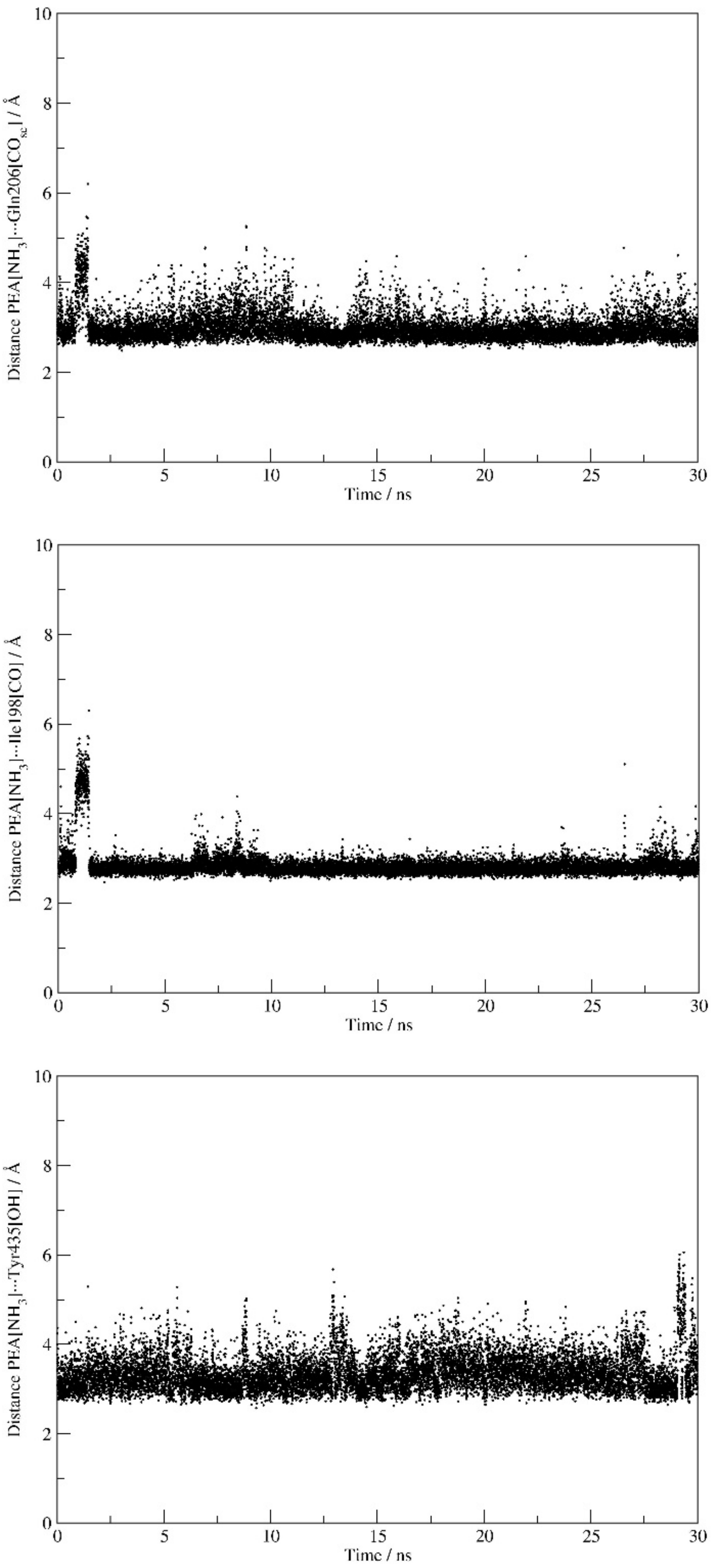


**Figure S8.** Evolution of distances between the protonated amine N-atom in **PEA** and (i) the side chain amide O-atom of Gln206 in **MAO B** (top), (ii) the backbone amide O-atom of Ile198 in **MAO B** (middle), and (iii) the side chain hydroxy O-atom from Tyr435 in **MAO B** (bottom), all in the **native MAO B** during MD simulations.

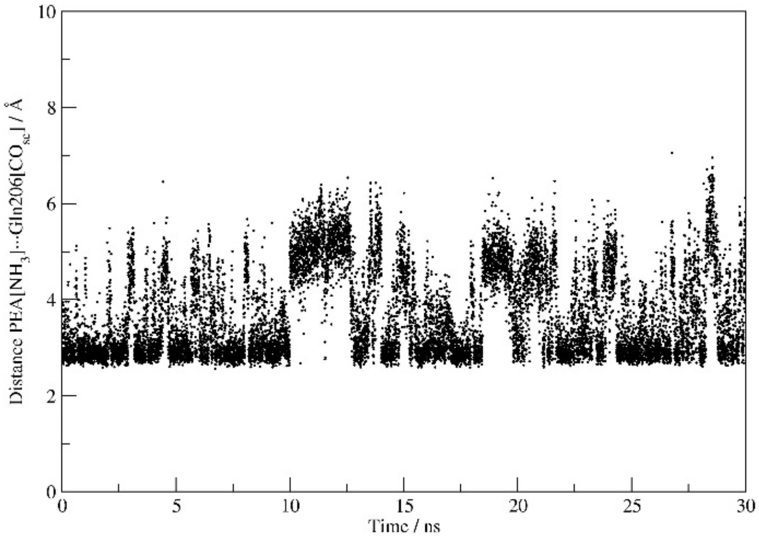


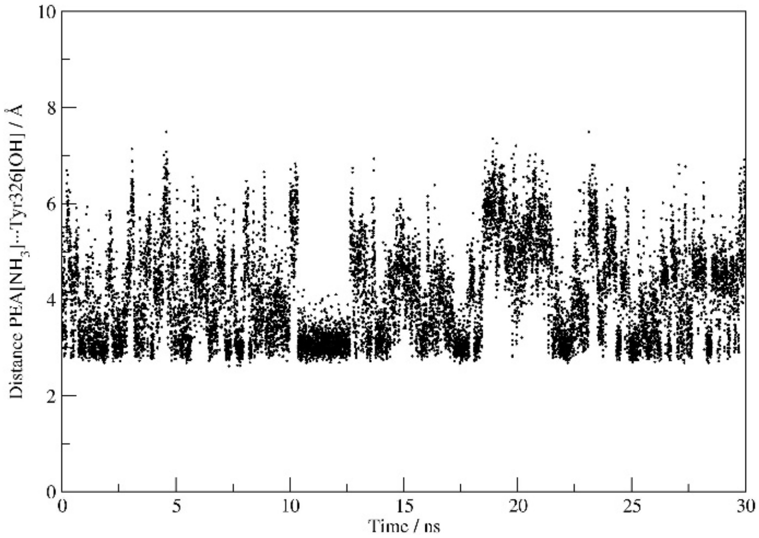


**Figure S9.** Evolution of distances between the protonated amine N-atom in **PEA** and (i) the side chain amide O-atom of Gln206 in **MAO B** (top), and (ii) the side chain hydroxy O-atom of Tyr326 in **MAO B** (bottom), all in the **MAO B···WT complex** during MD simulations. The results pertain to the **MAO B** subunit directly interacting with the matching spike protein.

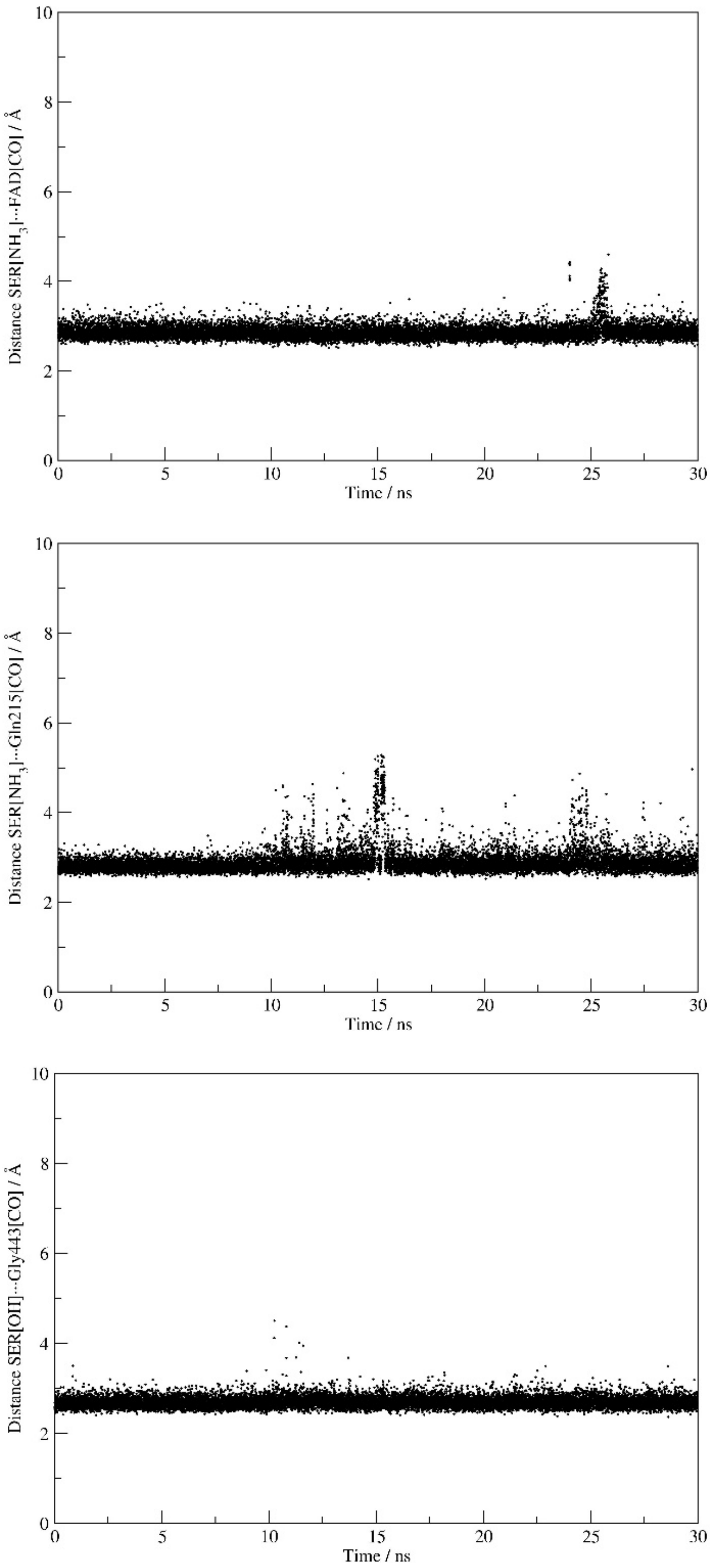


**Figure S10.** Evolution of distances between the protonated amine N-atom in **SER** and (i) the carbonyl O-atom of the FAD cofactor in **MAO A** (top) and (ii) the side chain amide O-atom of Gln215 in **MAO A** (middle), and between the hydroxy O-atom in **SER** and (iii) the backbone amide O-atom of Gly443 in **MAO A** (bottom), all in the **native MAO A** during MD simulations.

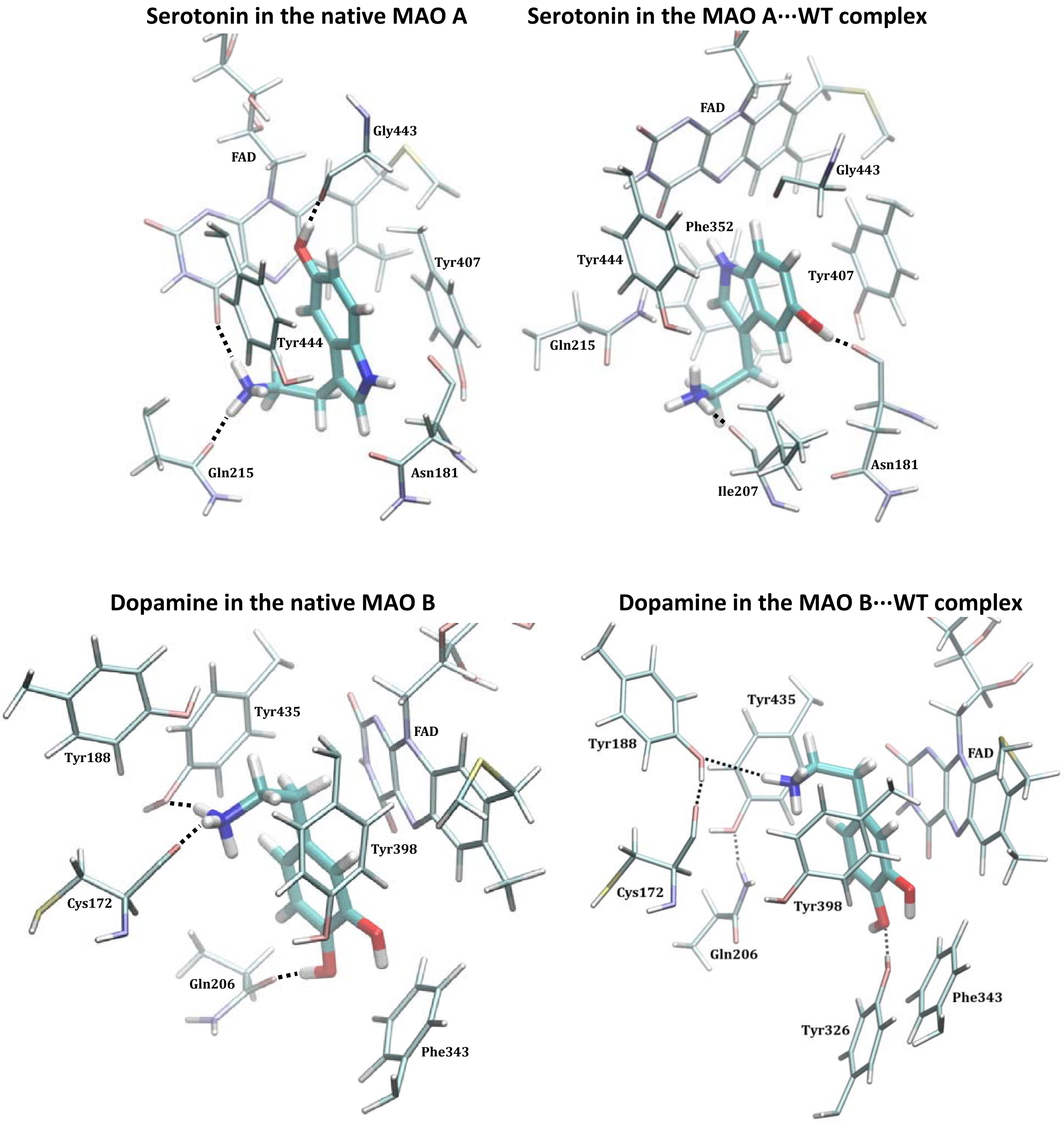


**Figure S11.** Binding positions of **SER** and **DOP** within the active sites of **native MAO A** and **MAO B** enzymes, and following their complex formation with the **WT** strain as obtained from MD simulations. The results for the **MAO B**···**WT** complex pertain to the **MAO B** subunit directly interacting with the matching spike protein.

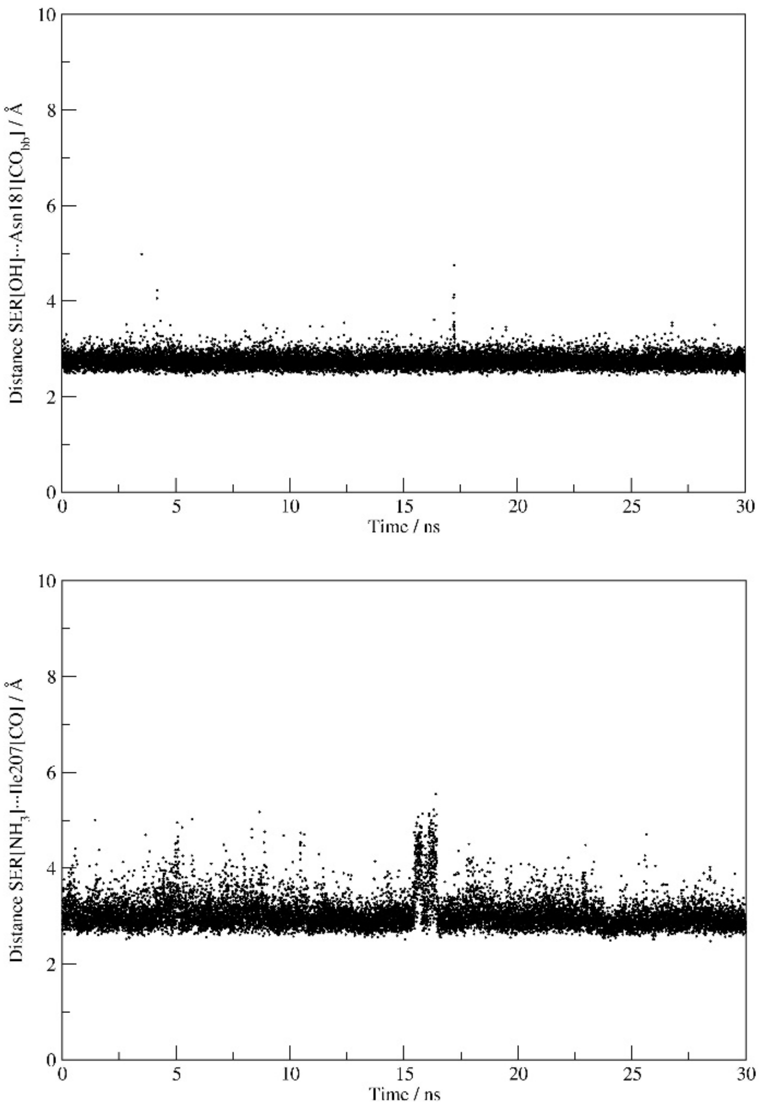


**Figure S12.** Evolution of distances between the hydroxy O-atom in **SER** and the backbone amide O-atom of Asn181 in **MAO A** (top), and the protonated amine N-atom in **SER** and the backbone amide O-atom of Ile207 in **MAO A** (bottom), all in the **MAO A···WT** complex during MD simulations.

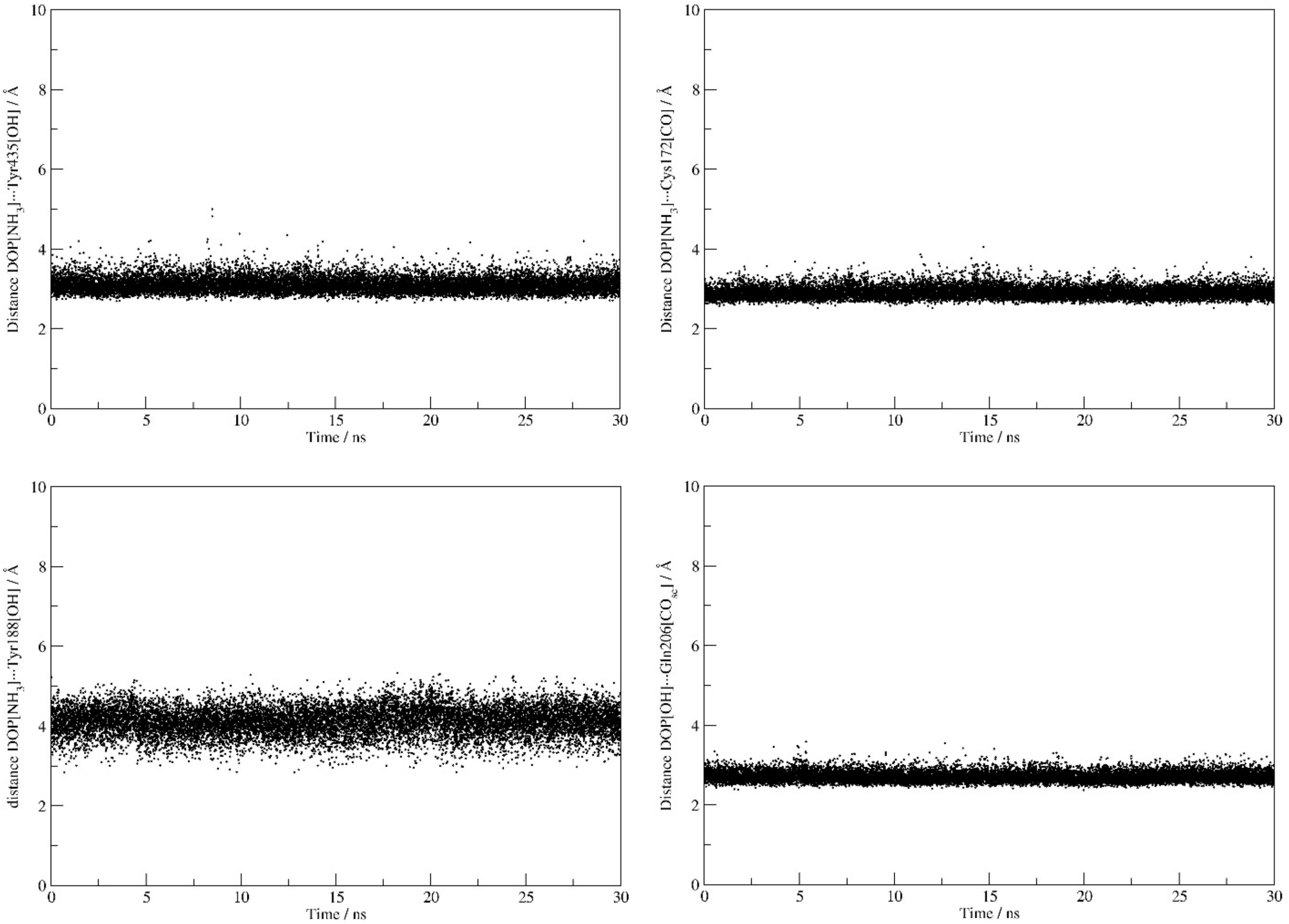

**Figure S13.** Evolution of distances between the protonated amine N-atom in **DOP** and (i) the side chain hydroxy O-atom of Tyr435 in **MAO B** (top row, left), (ii) the backbone amine O-atom of Cys172 in **MAO B** (top row, right), and (iii) the side chain hydroxy O-atom of Tyr188 in **MAO B** (bottom row, left), and between the hydroxy O-atom in **DOP** and the side chain amide O-atom of Gln206 in **MAO B** (bottom row, right), all in the **native MAO B** during MD simulations.

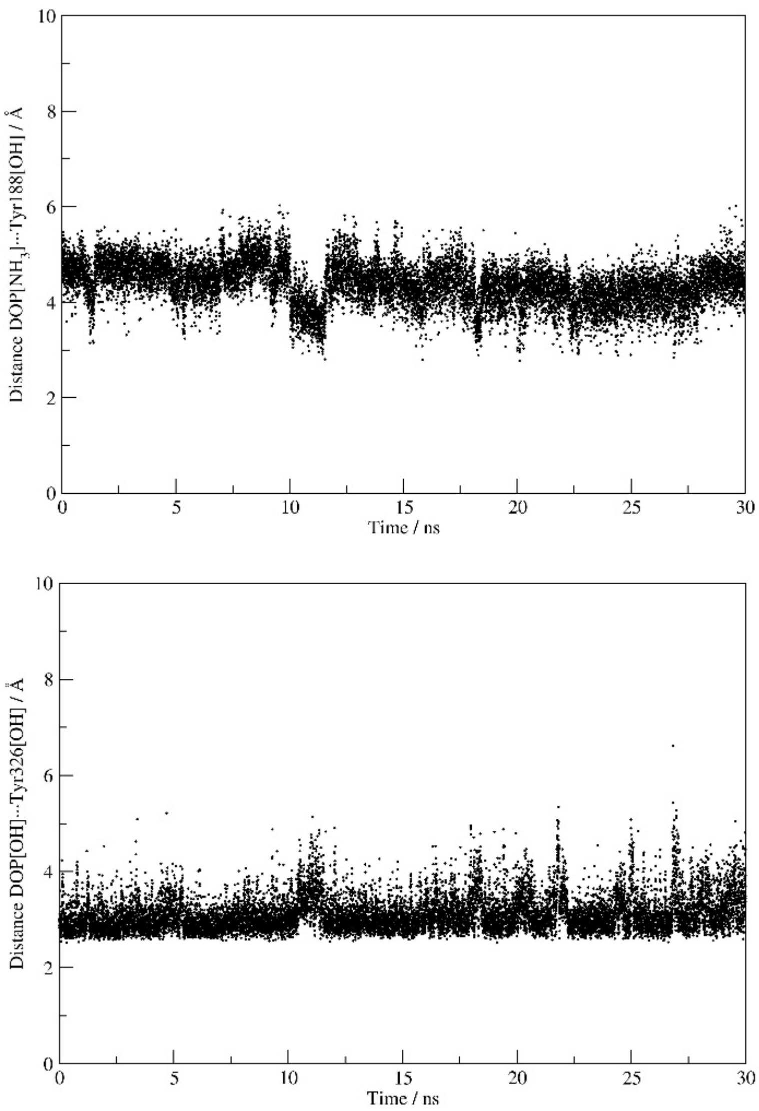


**Figure S14.** Evolution of distances between the protonated amine N-atom in **DOP** and the side chain hydroxy O-atom of Tyr188 in **MAO B** (top), and between the hydroxy O-atom in **DOP** and the side chain hydroxy O-atom of Tyr326 in **MAO B** (bottom), all in the **MAO B···WT** complex during MD simulations. The results pertain to the **MAO B** subunit directly interacting with the matching spike protein.

**Table S4.** Calculated binding free energies ($\Delta G_{BIND}$) from molecular dynamic trajectories using the MM-GBSA approach for **MAO A** substrates, and their decomposition on a *per-residue* basis in the **MAO A** enzyme (in kcal mol$^{-1}$).[a]

| **MAO A** | | | | | | | |
|---|---|---|---|---|---|---|---|
| **SEROTONIN** | | | | **PHENYLETHYLAMINE** | | | |
| **$\Delta G_{BIND}$ (kcal/mol)** | −20.1 | −15.5 | −23.0 | **$\Delta G_{BIND}$ (kcal/mol)** | −16.8 | −17.0 | −15.8 |
| **Residue** | **MAO A** | **MAO A···WT** | **MAO A···SA** | **Residue** | **MAO A** | **MAO A···WT** | **MAO A···SA** |
| FAD | −6.65 | −2.70 | −3.04 | Gln215 | −4.86 | −3.91 | −3.40 |
| Tyr407 | −2.52 | −2.24 | −2.18 | FAD | −2.58 | −3.28 | −1.36 |
| Tyr444 | −2.32 | −1.30 | −5.42 | Tyr407 | −1.98 | −2.23 | −1.00 |
| Gln215 | −1.98 | −0.44 | −1.85 | Phe352 | −1.06 | −0.89 | −1.05 |
| Gly443 | −1.76 | −0.12 | 0.19 | Tyr69 | −0.83 | −0.79 | −0.45 |
| Asn181 | −1.06 | −4.39 | −3.81 | Phe208 | −0.65 | −0.82 | −0.50 |
| Tyr69 | −0.85 | −0.16 | −0.35 | Ile335 | −0.41 | −0.45 | −1.19 |
| Phe208 | −0.75 | −0.05 | −0.79 | Tyr444 | −0.40 | −0.61 | −0.72 |
| Phe352 | −0.57 | −1.08 | −0.25 | Ile180 | −0.40 | −0.30 | −0.55 |
| Val182 | −0.48 | −0.55 | −0.48 | Leu337 | −0.39 | −0.31 | −0.50 |
| Ile207 | −0.36 | −1.54 | −0.33 | Val210 | −0.19 | 0.00 | −0.33 |
| Thr408 | −0.32 | −0.28 | −0.05 | Trp441 | −0.14 | −0.10 | −0.14 |
| | | Ile335<br>−0.30 | Tyr197<br>−3.09 | | | Glu436<br>−0.15 | Asn181<br>−0.22 |
| | | Ile180<br>−0.19 | Ile180<br>−0.90 | | | Ile207<br>−0.15 | |
| | | | Glu436<br>−0.30 | | | | |

[a]Residues are selected to list those most dominant for the binding of each substrate to the **native MAO A**, while residues with an increased importance in the complexes with the **WT** and **SA** strains are additionally presented in shading.

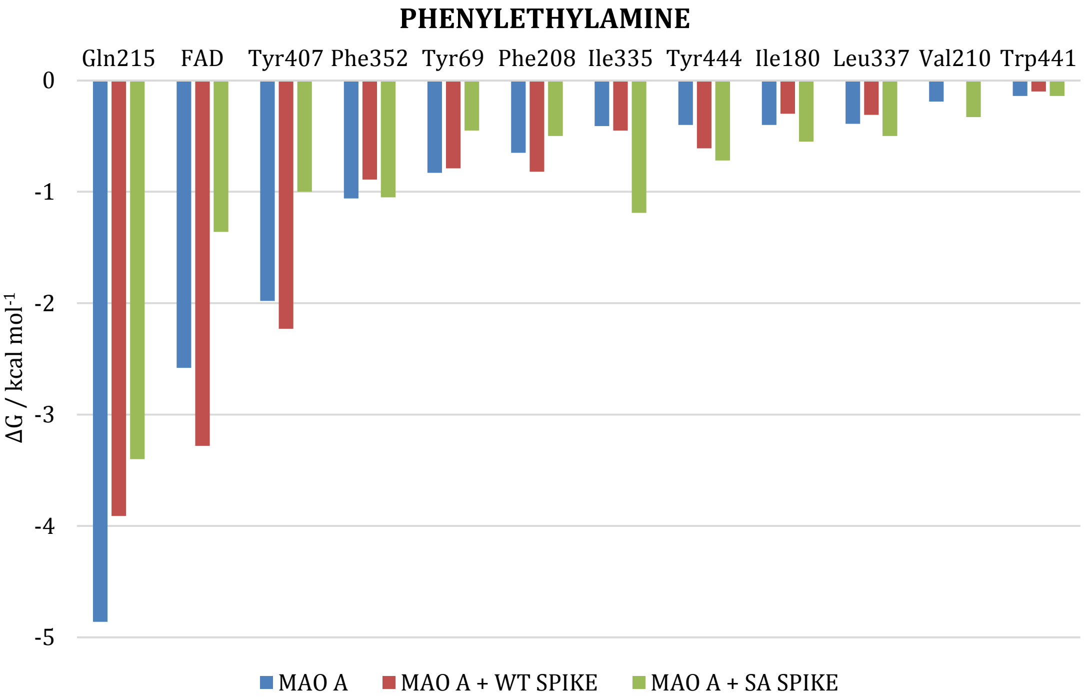


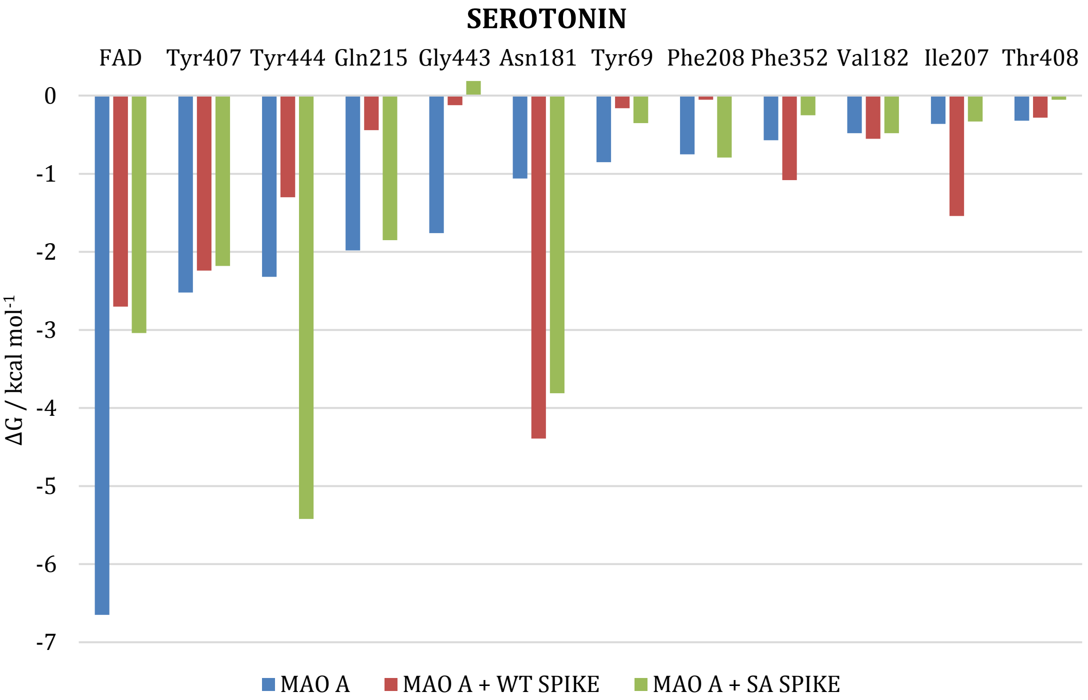


**Figure S15.** Graphical representations of the most dominant residues governing the binding of **PEA** (top) and **SER** (bottom) to the **native MAO A** enzyme, and their changes induced by complex formation with the **WT** and **SA** strains, as obtained from the MM-GBSA analysis.

**Table S5.** Calculated binding free energies ($\Delta G_{BIND}$) from molecular dynamic trajectories using the MM-GBSA approach for **MAO B** substrates, and their decomposition on a *per-residue* basis in the **MAO B** enzyme (in kcal $mol^{-1}$).[a]

| MAO B | | | | | | | |
|---|---|---|---|---|---|---|---|
| DOPAMINE | | | | PHENYLETHYLAMINE | | | |
| **$\Delta G_{BIND}$ (kcal/mol)** | −20.7 | −15.4 | −23.0 | **$\Delta G_{BIND}$ (kcal/mol)** | −12.0 | −9.4 | −14.8 |
| **Residue** | **MAO B** | **MAO B···WT** | **MAO B···SA** | **Residue** | **MAO B** | **MAO B···WT** | **MAO B···SA** |
| Tyr435 | −5.27 | −3.65 | −2.21 | Gln206 | −3.28 | −0.63 | −0.24 |
| FAD | −3.63 | −3.98 | −0.99 | Ile198 | −2.95 | −0.13 | −0.11 |
| Gln206 | −2.37 | −0.17 | −0.75 | Tyr435 | −0.97 | −1.64 | −4.55 |
| Cys172 | −2.13 | −0.08 | −1.51 | Tyr326 | −0.73 | −0.77 | −0.51 |
| Tyr398 | −2.07 | −4.64 | −2.40 | Tyr398 | −0.72 | −1.40 | −2.11 |
| Phe343 | −1.25 | −1.36 | −0.12 | Tyr60 | −0.60 | −0.22 | −0.09 |
| Tyr188 | −0.96 | −0.50 | −3.48 | Ile199 | −0.60 | −0.06 | −1.08 |
| Tyr60 | −0.86 | −0.63 | −0.07 | Cys172 | −0.57 | −0.09 | −2.23 |
| Ile199 | −0.76 | −0.25 | −1.52 | FAD | −0.52 | −4.54 | −0.71 |
| Tyr326 | −0.57 | −0.56 | −0.86 | Leu171 | −0.52 | 0.03 | −0.62 |
| Val173 | −0.53 | −0.02 | −0.33 | Leu328 | −0.38 | −0.06 | −0.02 |
| Glu427 | −0.22 | −0.21 | −0.44 | Ser200 | −0.25 | 0.00 | −0.09 |
| | | Gly58 | Ser433 | | | Phe343 | Tyr188 |
| | | −0.29 | −3.60 | | | −0.44 | −1.57 |
| | | Asp55 | Leu171 | | | Asp55 | Val173 |
| | | −0.22 | −2.38 | | | −0.23 | −0.51 |
| | | Thr399 | Cys192 | | | Thr399 | Glu427 |
| | | −0.20 | −2.15 | | | −0.19 | −0.24 |
| | | | Ile198 | | | Gly58 | Cys192 |
| | | | −0.64 | | | −0.17 | −0.21 |
| | | | Gln191 | | | Gly434 | Thr174 |
| | | | −0.26 | | | −0.16 | −0.18 |
| | | | Trp184 | | | Val173 | |
| | | | −0.25 | | | −0.11 | |
| | | | Gly434 | | | | |
| | | | −0.25 | | | | |

[a]Residues are selected to list those most dominant for the binding of each substrate to the **native MAO B**, while residues with an increased importance in the complexes with the **WT** and **SA** strains are additionally presented in shading. The results for the **WT/SA**···**MAO B** complexes pertain to the **MAO B** subunit directly interacting with the matching spike protein.

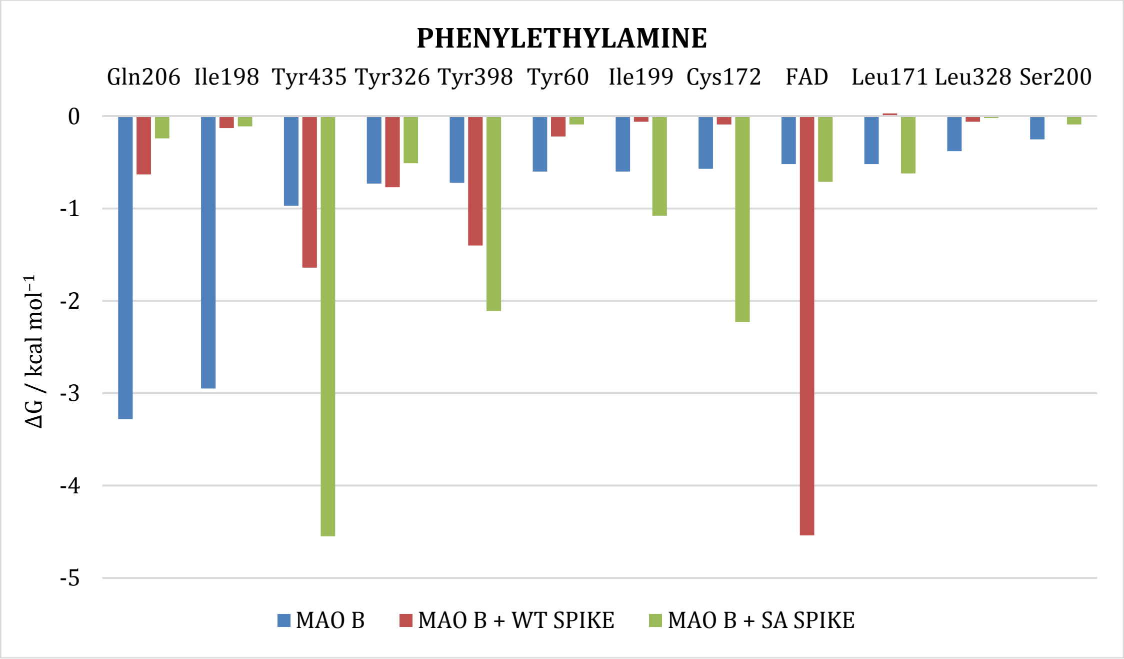


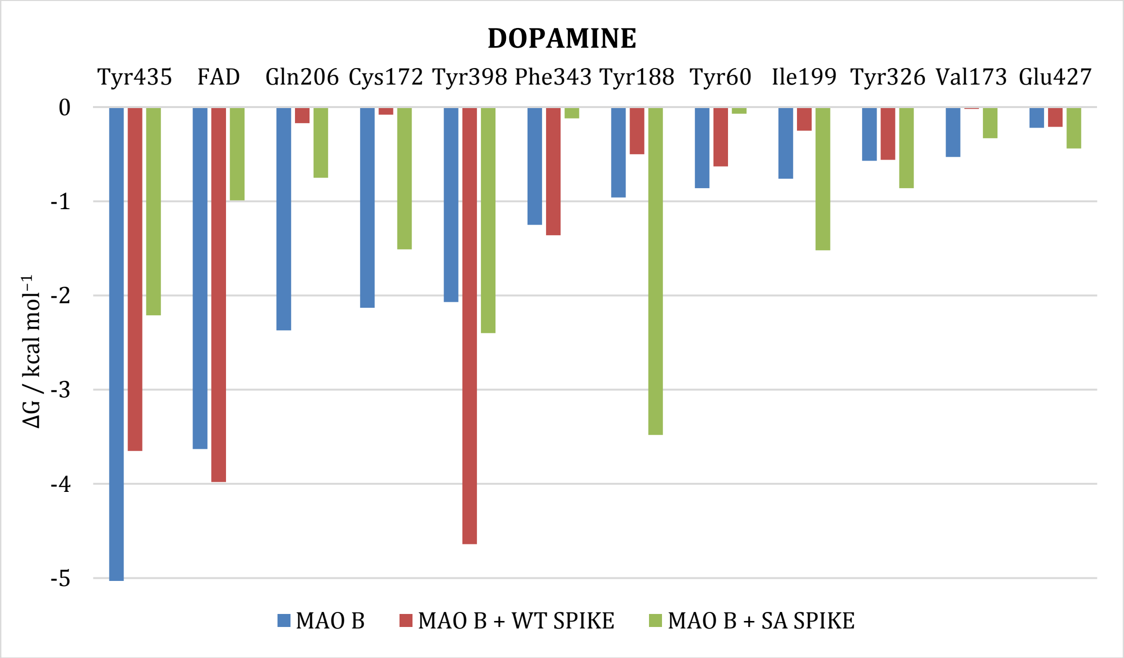


**Figure S16.** Graphical representations of the most dominant residues governing the binding of **PEA** (top) and **DOP** (bottom) to the **native MAO B** enzyme, and their changes induced by complex formation with the **WT** and **SA** strains, as obtained from the MM-GBSA analysis. The results for the **WT/SA···MAO B** complexes pertain to the **MAO B** subunit directly interacting with the matching spike protein.

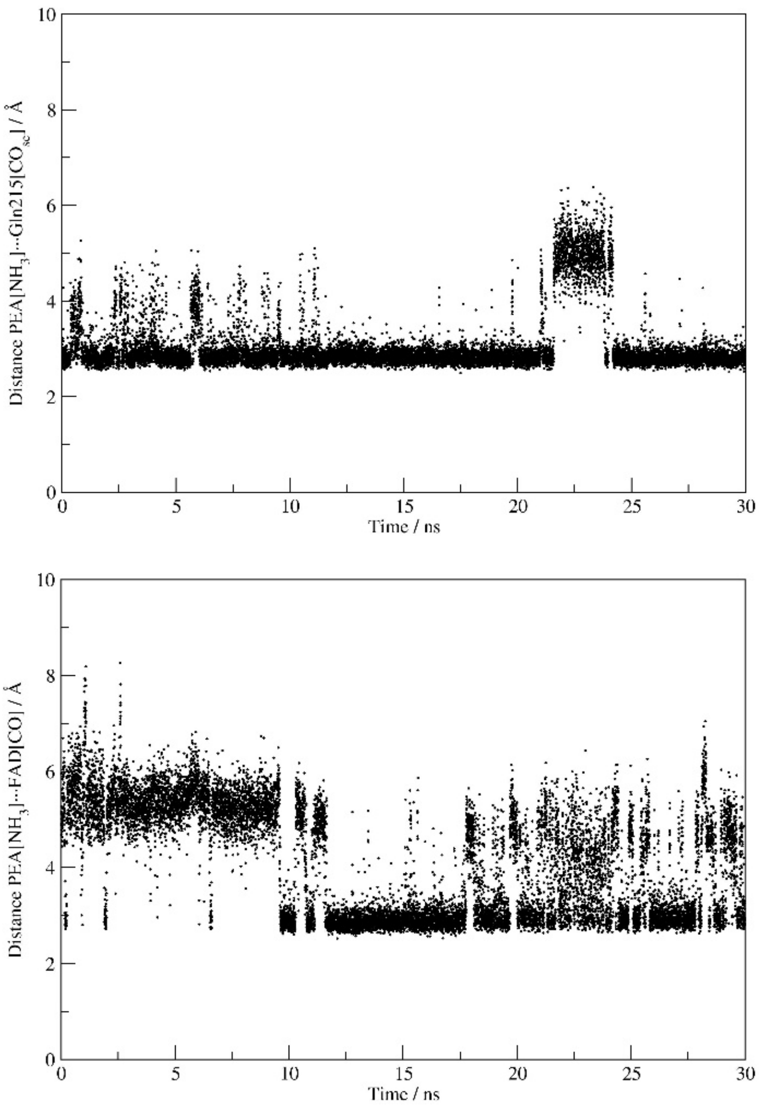


**Figure S17.** Evolution of distances between the protonated amine N-atom in **PEA** and (i) the side chain amide O-atom of Gln215 in **MAO A** (top), and (ii) the carbonyl O-atom of the FAD cofactor in **MAO A** (bottom), all in the **MAO A···SA** complex during MD simulations.

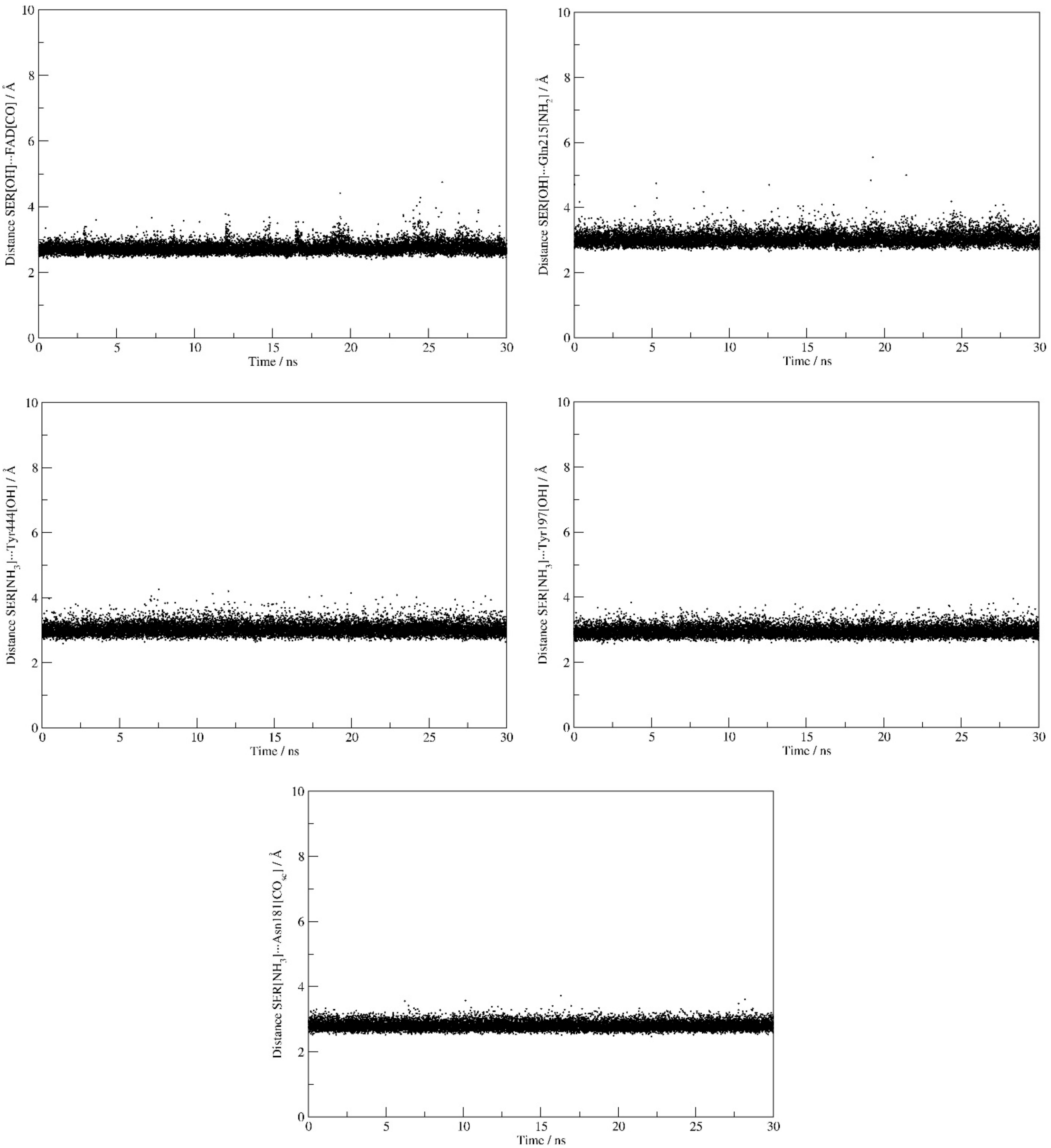


**Figure S18.** Evolution of distances between the hydroxy O-atom in **SER** and (i) the carbonyl O-atom of the FAD cofactor in **MAO A** (top row, left) and (ii) the side chain amide N-atom of Gln215 in **MAO A** (top row, right), and between the protonated amine N-atom in **SER** and (iii) the side chain hydroxy O-atom of Tyr444 in **MAO A** (middle row, left), (iv) the side chain hydroxy O-atom of Tyr197 in **MAO A** (middle row, right), and (v) the side chain amide O-atom of Asn181 in **MAO A** (bottom row), all in the **MAO A···SA** complex during MD simulations.

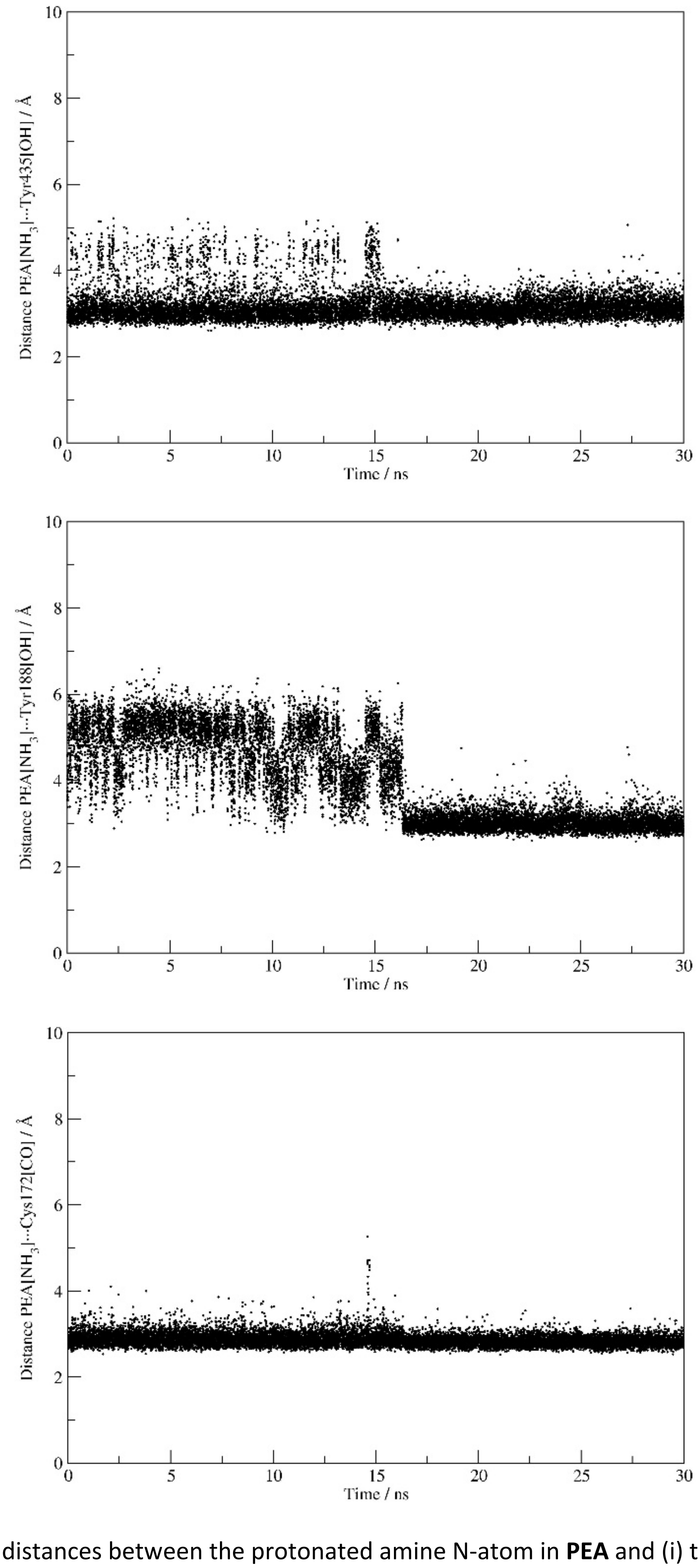


**Figure S19.** Evolution of distances between the protonated amine N-atom in **PEA** and (i) the side chain hydroxy O-atom of Tyr435 in **MAO B** (top), (ii) the side chain hydroxy O-atom of Tyr188 in **MAO B** (middle) and (iii) the backbone amide O-atom of Cys172 in **MAO B** (bottom), all in the **MAO B···SA** complex during MD simulations. The results pertain to the **MAO B** subunit directly interacting with the matching spike protein.

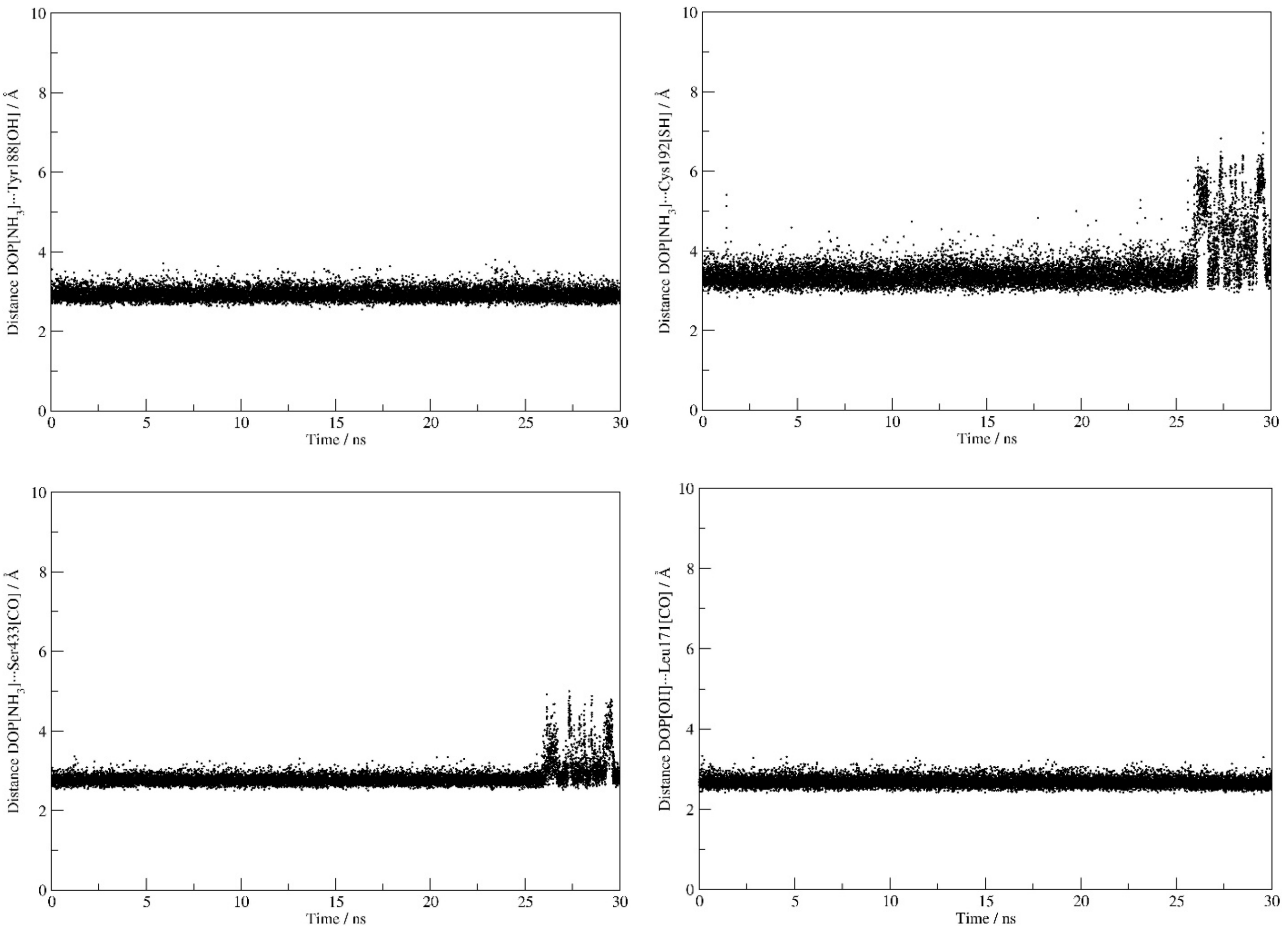


**Figure S20.** Evolution of distances between the protonated amine N-atom in **DOP** and (i) the side chain hydroxy O-atom of Tyr188 in **MAO B** (top row, left), (ii) the side chain thiol S-atom of Cys192 in **MAO B** (top row, right), (iii) the backbone amide O-atom of Ser433 in **MAO B** (bottom row, left), and between the hydroxy O-atom in **DOP** and (iv) the backbone amide O-atom in Leu171 (bottom row, right), all in the **MAO B···SA** complex during MD simulations. The results pertain to the **MAO B** subunit directly interacting with the matching spike protein.

### COMPUTATIONAL DETAILS

#### Preparation of MAO proteins within a membrane

The X-ray crystal structures of human MAO A in a complex with harmine (PDB ID: 2Z5X)[S2] and human MAO B in a complex with *N*-methyl-1(*R*)-aminoindan (PDB ID: 2C67)[S3] were used as starting points for the analysis. The co-crystallized ligands, water molecules and ions were removed, while the protonation states of protein ionizable residues were set according to PROPKA3.1 server predictions[S4] and by inspecting hydrogen bonding networks in their closest vicinity. The missing hydrogen atoms were added using the *tleap* module in AmberTools16.[S5] Structures of MAO enzymes obtained in this way were then inserted into a lipid bilayer consisting of zwitterionic, electrically neutral 1,2-dioleoyl-*sn*-glycero-3-phosphocholine (DOPC) using the CHARMM-GUI web-server.[S6] For MAO A, we used 75 molecules in both layers, while for the dimeric MAO B, the number of DOPC molecules in each layer was 120. The position of both MAO enzymes in the membrane was determined using the OPM database through the PPM web server.[S7] Each complex was solvated using explicit TIP3P water molecules above and below the membrane, while making sure that the entire protein structure is located within the solvent box. Also, an overall concentration of 0.15 M of $K^+$ and $Cl^-$ ions was distributed around the system using the Monte-Carlo methodology within CHARMM-GUI to maintain the net zero charge in all systems. The obtained structures were parameterized using the AMBER ff14SB and LIPID17 force fields for the protein and lipid components, respectively, while the parameters for the MAO's FAD cofactor in its oxidized form were taken from our earlier study.[S8] These were submitted to geometry optimization in the AMBER16 program package,[S5] employing periodic boundary conditions in all directions. In all instances, optimized systems were gradually heated from 0 to 300 K and equilibrated during 30 ps using NVT conditions, followed by a productive and unconstrained MD simulation of 300 ns, employing a time step of 2 fs at a constant pressure (1 atm) and temperature (300 K), the latter held constant using a Langevin thermostat with a collision frequency of 1 $ps^{-1}$. The long-range electrostatic interactions were calculated employing the Particle Mesh Ewald method,[S9] and were updated in every second step, while the non-bonded interactions were truncated at 11.0 Å. Such a setup was identically applied to all MD simulations employed in this work. Following the MD simulations, we extracted representative structures of both MAO isoforms immersed in the membrane (Figure S21) and used them for the subsequent docking studies and molecular dynamic simulations.

#### Preparation of the ACE2 receptor and the spike protein S1 receptor-binding domain subunit from the SARS-CoV-2 wild type and its South African B.1.351 variant

The X-ray crystal structure of the h-ACE2 receptor in complex with the spike protein from the SARS-CoV-2 wild type (PDB ID: 6M0J)[S10] was used to extract the geometry of the receptor and the S1 receptor-binding domain from the WT strain, while an analogous structure corresponding to the South African mutant variant B.1.351 was taken from the PDB ID: 7LYN.[S11] The co-crystallized ligands, water molecules and ions were removed, and the resulting structures were separately submitted to docking studies in order to verify the procedure in terms of the obtained binding positions and their comparison with the original crystal structures.

**Protein-protein docking studies and subsequent molecular dynamic simulations**

Protein-protein docking was performed to determine relevant binding poses for the spike protein in the complex with the ACE2 receptor in one case, and with both MAO isoforms in the other. In all instances, the structures of the obtained complexes were used for subsequent analysis, while the elucidated spike protein···ACE2 binding conformations were additionally employed to validate the docking procedure through the comparison with the mentioned crystal structures. At this point, it is worth to reiterate that MAO enzymes were, in all cases, submitted to docking analysis as immersed in the DOPC membrane determined earlier, which prevented the spike protein to artificially bind areas around the MAO enzymes that are otherwise inaccessible due to the membrane presence. We came to this decision after initially performing docking analysis without the explicit membrane, which resulted in WT/SA···MAO complexes having the spike protein bind the membrane-bound areas in MAO enzymes as the most favorable positions.

Docking studies were performed using the HDOCK web-server[S12] with default parameters for all docking runs. HDOCK server uses an iterative knowledge-based scoring function referred to as an ITScore-PP[S13] to rank the top 10 binding poses. In the case of the WT/SA···ACE2 complexes, only the most stable binding interaction was considered, after realizing that it matches the corresponding crystal structure. For the WT···MAO interactions, all ten binding poses were clustered into three distinct positions for the monomeric MAO A, while for a more complex dimeric MAO B, all ten poses appeared to bind different MAO regions, and were all considered for further analysis. Analogously, in the case of the SA···MAO interactions, with both MAO isoforms we recognized five discrete binding positions, which were further considered as such.

All of the obtained structures, one for the WT···ACE2 and SA···ACE2 complexes, three for the WT···MAO A complex, five for the SA···MAO A and SA···MAO B complexes, and ten for the WT···MAO B complexes, were solvated in a truncated octahedral box of TIP3P water molecules spanning a 10 Å-thick buffer, and submitted to molecular dynamic simulations in AMBER16 using the same setup as already described. The obtained MD trajectories, and thereof considered protein-protein complexes, were evaluated and ranked by the protein-protein binding free energies, $\Delta G_{\mathrm{BIND}}$, using the established MM-GBSA protocol [S14,S15] available in AmberTools16,[S5] and in line with our earlier reports.[S16,S17] MM-GBSA is widely used for calculating the relative binding free energies from snapshots of the MD trajectory with an estimated standard error of 1–3 kcal mol$^{-1}$.[S14] For that purpose, 3.000 snapshots, collected from the last 30 ns of the corresponding MD trajectories, were used. The calculated MM-GBSA binding free energies were decomposed into specific residue contributions on a *per-residue* basis according to an established procedure.[S18,S19] This protocol evaluates $\Delta G_{\mathrm{BIND}}$ contributions from each amino acid residue and identifies the nature of the energy change in terms of interaction and solvation energies or entropic contributions. The trajectories corresponding to the most exergonic binding were used to extract the representative structures of each complex, which were used in the subsequent analysis.

**Docking of the neurotransmitter substrates to the active site of native MAO enzymes and MAO enzymes complexed with the WT/SA spike protein and subsequent molecular dynamic simulations**

To evaluate changes in the activity of the MAO enzymes following the complex formation with the WT/SA spike protein, we performed docking studies of phenylethylamine (**PEA**) to both MAO isoforms, and their more specific substrates, serotonin (**SER**) to MAO A, and dopamine (**DOP**) to MAO B. In all cases, the docking analysis was used separately for native MAO enzymes and their complexes with the spike protein from both the WT and the SA strains. All three substrates were considered with a protonated amino group, since their aqueous phase p$K_a$ values are found to be between 9–10 in all cases,[S20] while earlier experimental[S21] and computational[S22] studies confirmed that MAO substrates bind within the active site as monocations.

Docking was performed using the AutoDock Vina program,[23] which uses dispersion, hydrogen bonds, electrostatic, and desolvation components to determine the most probable complex conformation. Structures of native MAO enzymes and MAO enzymes complexed with the WT/SA spike protein were taken as representative structures from the aforementioned molecular dynamics simulations. For docking purposes, all water molecules were omitted from the analysis, hydrogen atoms were added were necessary, and all Lys, Arg, and His side chains were protonated, while all Asp and Glu side chains were deprotonated, and both amino and carboxyl ends were charged using the UCSF Chimera 1.14. program (University of California, USA),[S24] which was also used for results visualization and interpretation. For all docking attempts, a grid map of size 40 × 40 × 40 Å was generated by the AutoGrid program[S25] and centered at the FAD cofactor centre of mass coordinates (in case of MAO B, docking to both subunits was performed). The receptor molecule was regarded as rigid, except the FAD cofactor and active site residues, while all single bonds within each ligand were allowed to rotate freely. All studies were run with both exhaustiveness and number of modes set to 100.

After docking protocol identified the most stable binding positions for all substrates within each considered biomolecular system, the obtained structures were submitted to the MD simulations using the same setup as already described. To parameterize the amine substrates for the MD simulations, their geometries were optimized in the Gaussian16 program,[S26] employing the M06–2X/6–31+G(d) level of theory and implicit SMD water solvation, while their RESP charges were calculated using the HF/6–31G(d) model to be consistent with the subsequently employed AMBER GAFF force field. All MD simulations for this part of the work were done in quadruplicates, and all trajectories were submitted to the MM–GBSA analysis to calculate the matching substrate binding affinities, $\Delta G_{BIND}$. In each case, the trajectory corresponding to the substrate binding affinity closest to the average of the mentioned four $\Delta G_{BIND}$ values was used for further analysis.

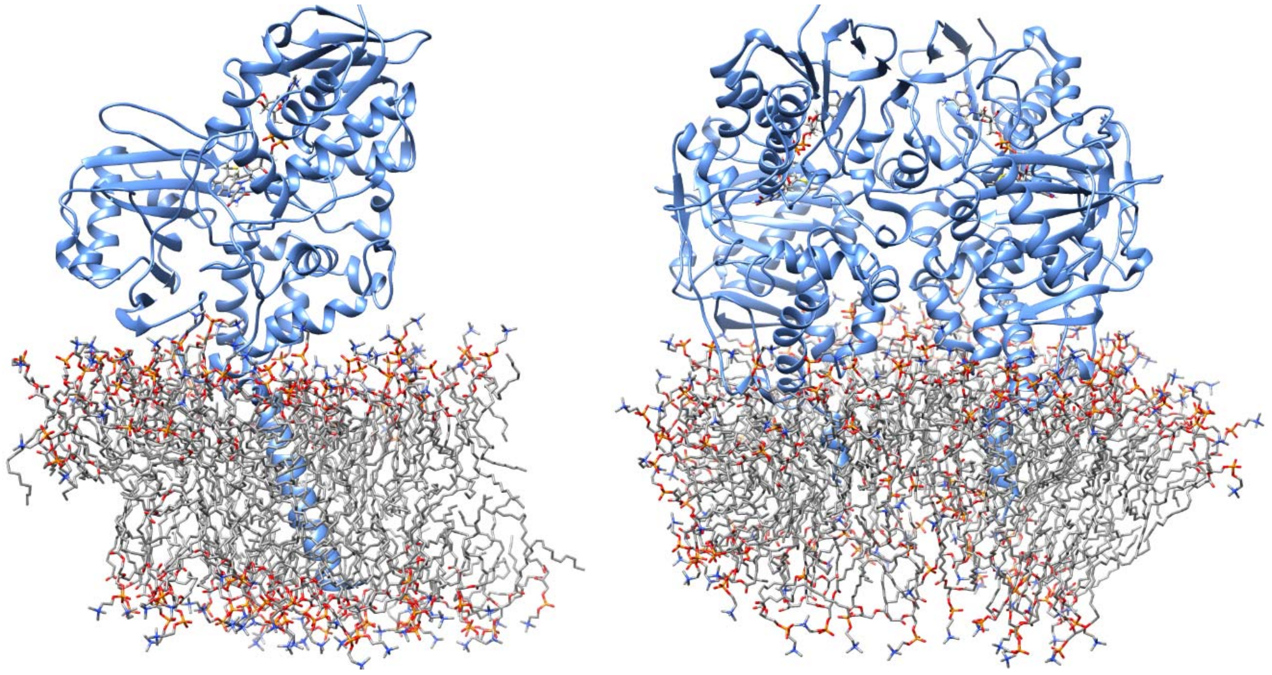

**Figure S21.** Equilibrated structures of MAO A (left) and MAO (B) immersed in a bilayer DOPC membrane following MD simulations. These structures were subsequently employed in all docking studies and molecular dynamic simulations, with the idea to prevent the docked spike protein and MAO substrates to artificially reside in the areas occupied by the membrane.

### REFERENCES


[S1] M. Ramanathan, I. D. Ferguson, W. Miao, P. A. Khavarib. SARS-CoV-2 B.1.1.7 and B.1.351 spike variants bind human ACE2 with increased affinity. *Lancet Infect Dis*, 2021, 21, 1070.

[S2] S.-Y. Son, J. Ma, Y. Kondou, M. Yoshimura, E. Yamashita, T. Tsukihara. Structure of human monoamine oxidase A at 2.2-A resolution: the control of opening the entry for substrates/inhibitors. *Proc Natl Acad Sci U S A*, 2008, 105, 5739–5744.

[S3] C. Binda, F. Hubálek, M. Li, Y. Herzig, J. Sterling, D. E. Edmondson, A. Mattevi. Binding of rasagiline-related inhibitors to human monoamine oxidases: a kinetic and crystallographic analysis. *J Med Chem*, 2005, 48, 8148–8154.

[S4] M. H. M. Olsson, C. R. Søndergaard, M. Rostkowski, J. H. Jensen. PROPKA3: Consistent treatment of internal and surface residues in empirical p$K_a$ predictions. *J Chem Theory Comput* 2011, 7, 525–537.

[S5] D. A. Case, R. M. Betz, D. S. Cerutti, T. E. Cheatham, III, T. A. Darden, R. E. Duke, T. J. Giese, H. Gohlke, A. W. Goetz, N. Homeyer, S. Izadi, P. Janowski, J. Kaus, A. Kovalenko, T. S. Lee, S. LeGrand, P. Li, C. Lin, T. Luchko, R. Luo, B. Madej, D. Mermelstein, K. M. Merz, G. Monard, H. Nguyen, H. T. Nguyen, I. Omelyan, A. Onufriev, D. R. Roe, A. Roitberg, C. Sagui, C. L. Simmerling, W. M. Botello-Smith, J. Swails, R. C. Walker, J. Wang, R. M. Wolf, X. Wu, L. Xiao, P. A. Kollman. *AMBER 2016*, 2016, University of California, San Francisco.

[S6] S. Jo, T. Kim, V. G. Iyer, W. Im. CHARMM-GUI: a web-based graphical user interface for CHARMM. *J Comput Chem*, 2008, 29, 1859–1865.

[S7] M. A. Lomize, I. D. Pogozheva, H. Joo, H. I. Mosberg, A. L. Lomize. OPM database and PPM web server: resources for positioning of proteins in membranes. *Nucleic Acids Res*, 2012, 40, D370–D376.

[S8] T. Tandarić, R. Vianello. Computational insight into the mechanism of the irreversible inhibition of monoamine oxidase enzymes by the anti-parkinsonian propargylamine inhibitors rasagiline and selegiline. *ACS Chem Neurosci*, 2019, 10, 3532–3542.

[S9] T. Darden, D. York, L. Pedersen. Particle Mesh Ewald: An $N$·log($N$) Method for Ewald Sums in Large Systems. *J Chem Phys*, 1993, 98, 10089–10092.

[S10] J. Lan, J. Ge, J. Yu, S. Shan, H. Zhou, S. Fan, Q. Zhang, X. Shi, Q. Wang, L. Zhang, X. Wang. Structure of the SARS-CoV-2 spike receptor-binding domain bound to the ACE2 receptor. *Nature*, 2020, 581, 215–220.

[S11] S. M.-C. Gobeil, K. Janowska, S. McDowell, K. Mansouri, R. Parks, V. Stalls, M. F. Kopp, K. Manne, D. Li, K. Wiehe, K. O. Saunders, R. J. Edwards, B. Korber, B. F. Haynes, R. Henderson, P. Acharya. Effect of natural mutations of SARS-CoV-2 on spike structure, conformation, and antigenicity. *Science*, 2021, 373, eabi6226.

[S12] Y. Yan, H. Tao, J. He, S.-Y. Huang. The HDOCK server for integrated protein–protein docking. *Nat Protoc*, 2020, 15, 1829–1852.

[S13] S.-Y. Huang, X. Zou. An iterative knowledge-based scoring function for protein-protein recognition. *Proteins*, 2008, 72, 557–579.

[S14] S. Genheden, U. Ryde. The MM/PBSA and MM/GBSA Methods to Estimate Ligand-Binding Affinities. *Expert Opin Drug Discov*, 2015, 10, 449–461.

[S15] T. Hou, J. Wang, Y. Li, W. Wang. Assessing the Performance of the MM/PBSA and MM/GBSA Methods. 1. The Accuracy of Binding Free Energy Calculations Based on Molecular Dynamics Simulations. *J Chem Inf Model*, 2011, 51, 69–82.

[S16] I. Perković, S. Raić-Malić, D. Fontinha, M. Prudêncio, L. Pessanha de Carvalho, J. Held, T. Tandarić, R. Vianello, B. Zorc, Z. Rajić. Harmicines – Harmine and Cinnamic Acid Hybrids as Novel Antiplasmodial Hits. *Eur J Med Chem*, 2020, 187, 111927.

[S17] L. Hok, J. Mavri, R. Vianello. The effect of deuteration on the $H_2$ receptor histamine binding profile: A computational insight into modified hydrogen bonding interactions. *Molecules*, 2020, 25, 6017.

[S18] H. Gohlke, C. Kiel, D. A. Case. Insights into Protein-Protein Binding by Binding Free Energy Calculation and Free Energy Decomposition for the Ras-Raf and Ras-RalGDS Complexes. *J Mol Biol*, 2003, 330, 891–913.

[S19] G. Rastelli, A. Del Rio, G. Degliesposti, M. Sgobba. Fast and Accurate Predictions of Binding Free Energies Using MM-PBSA and MM-GBSA. *J Comput Chem*, 2010, 31, 797–810.

[S20] S. Tshepelevitsh, A. Kütt, M. Lõkov, I. Kaljurand, J. Saame, A. Heering, P. Plieger, R. Vianello, I. Leito. On the Basicity of Organic Bases in Different Media. *Eur J Org Chem*, 2019, 6735–6748.

[S21] T. Z. E. Jones, D. Balsa, M. Unzeta, R. R. Ramsay. Variations in activity and inhibition with pH: the protonated amine is the substrate for monoamine oxidase, but uncharged inhibitors bind better. *J Neural Transm*, 2007, 114, 707.

[S22] R. Borštnar, M. Repič, S. C. L. Kamerlin, R. Vianello, J. Mavri. Computational study of the pKa values of potential catalytic residues in the active site of monoamine oxidase B. *J Chem Theory Comput*, 2012, 8, 3864–3870.

[S23] O. Trott, A. J. Olson. AutoDock Vina: Improving the speed and accuracy of docking with a new scoring function, efficient optimization, and multithreading. *J Comput Chem*, 2009, 31, 455–461.

[S24] E. F. Pettersen, T. D. Goddard, C. C. Huang, G. S. Couch, D. M. Greenblatt, E. C. Meng, T. E. Ferrin. UCSF Chimera – A visualization system for exploratory research and analysis. *J Comput Chem* 2004, 25, 1605–1612.

[S25] G. M. Morris, R. Huey, W. Lindstrom, M. F. Sanner, R. K. Belew, D. S. Goodsell, A. J. Olson. AutoDock4 and AutoDockTools4: Automated docking with selective receptor flexibility. *J Comput Chem* 2009, 30, 2785–2791.

[S26] Gaussian 16, Revision C.01. M. J. Frisch, G. W. Trucks, H. B. Schlegel, G. E. Scuseria, M. A. Robb, J. R. Cheeseman, G. Scalmani, V. Barone, G. A. Petersson, H. Nakatsuji, X. Li, M. Caricato, A. V. Marenich, J. Bloino, B. G. Janesko, R. Gomperts, B. Mennucci, H. P. Hratchian, J. V. Ortiz, A. F. Izmaylov, J. L. Sonnenberg, D. Williams-Young, F. Ding, F. Lipparini, F. Egidi, J. Goings, B. Peng, A. Petrone, T. Henderson, D. Ranasinghe, V. G. Zakrzewski, J. Gao, N. Rega, G. Zheng, W. Liang, M. Hada, M. Ehara, K. Toyota, R. Fukuda, J. Hasegawa, M. Ishida, T. Nakajima, Y. Honda, O. Kitao, H. Nakai, T. Vreven, K. Throssell, J. A. Montgomery, Jr., J. E. Peralta, F. Ogliaro, M. J. Bearpark, J. J. Heyd, E. N. Brothers, K. N. Kudin, V. N. Staroverov, T. A. Keith, R. Kobayashi, J. Normand, K. Raghavachari, A. P. Rendell, J. C. Burant, S. S. Iyengar, J. Tomasi, M. Cossi, J. M. Millam, M. Klene, C. Adamo, R. Cammi, J. W. Ochterski, R. L. Martin, K. Morokuma, O. Farkas, J. B. Foresman, D. J. Fox. *Gaussian*, Inc., 2016, Wallingford CT.
